# ZO-2 induces cytoplasmic retention of YAP by promoting a LATS1-ZO-2-YAP complex at tight junctions

**DOI:** 10.1101/355081

**Authors:** Olivia Xuan Liu, Lester Bocheng Lin, Tiweng Chew, Fumio Motegi, Boon Chuan Low

**Affiliations:** Department of Biological Sciences, National University of Singapore, Singapore 117543, Republic of Singapore; Mechanobiology Institute, National University of Singapore, Singapore 117411, Republic of Singapore; Temasek Life-sciences Laboratory, Singapore 117604, Republic of Singapore; University Scholars Programme, National University of Singapore, Singapore 138593, Republic of Singapore.

**Author notes:** These authors contributed equally to this work (Olivia Xuan Liu, Lester Bocheng Lin).

## Abstract

Contact inhibition of proliferation (CIP) is a key mechanism that transduces the cell density status of tissue and organs into a unique transcriptional program by translocating YAP between the nucleus and the cytoplasm. However, the nature of the cell density-dependent cues that regulate the YAP distribution remains unclear. Here, we present evidence that tight junctions serve as a platform that controls both distribution and activity of LATS1, a kinase that phosphorylates YAP. This CIP effect is mediated by the scaffold function of junctional protein, ZO-2, by promoting LATS1 interaction with YAP in the cytoplasm, and then targeting the tripartite complex to tight junctions. There, LATS1 is activated by angiomotin and NF2, thereby stimulating YAP phosphorylation and its cytoplasmic retention. Our findings delineate novel mechanisms governing CIP, in which ZO-2 utilizes the status of cell-cell cohesion to control the phosphorylation status and therefore inactivation of YAP by LATS1 in the cytoplasm.

## Introduction

The capacity of cells to regulate their proliferation in response to the extracellular environment is a fundamental requirement for the organization and maintenance of tissues and organs in multicellular organisms. Contact inhibition of proliferation (CIP) is a unique property of cellular communication that enables cells to stop growing after reaching a critical cell density (Edgar, 2006; McClatchey and Yap, 2012). Loss of CIP is a hallmark of many cancer cells, which have lost their ability to sense cell density and subsequently exhibit uncontrollable cell proliferation and tissue overgrowth (Edgar, 2006; McClatchey and Yap, 2012). The Hippo pathway has emerged as a conserved signalling pathway that controls cell proliferation in response to the extracellular environment (Gumbiner and Kim, 2014; Pan, 2010; Panciera et al., 2017; Yu et al., 2015; Zhao et al., 2007). The core signalling of the Hippo pathway consists of several protein kinases, which regulate the activity of the transcriptional co-activator, YAP, and its paralog, TAZ (Pan, 2010; Panciera et al., 2017; Yu et al., 2015). Both YAP and TAZ bind to transcriptional enhancer associate domain (TEAD) transcription factors, and coactivate the expression of proliferation-promoting genes (Galli et al., 2015; Huang et al., 2005; Vassilev et al., 2001; Wu et al., 2008; Zanconato et al., 2015). The MST-1 and MST-2 kinases trigger phosphorylation and activation of the LATS kinases (Callus et al., 2006; Chan et al., 2005; Wei et al., 2007; Wu et al., 2003), which in turn phosphorylates and inactivates YAP and TAZ by restricting them to the cytoplasm (Dong et al., 2007; Huang et al., 2005; Lei et al., 2008; Zhao et al., 2007). Hence, the activity of YAP and TAZ is primarily regulated by their subcellular distribution: YAP and TAZ in the cytoplasm are inactive, whereas those in the nucleus are active and stimulate cell proliferation. Several studies have shown that the subcellular localization of YAP and TAZ is affected by cell density; YAP and TAZ predominantly accumulates in the nucleus at low cell density and in the cytoplasm at high cell density (Kim et al., 2011; Schlegelmilch et al., 2011; Zhao et al., 2007). As cells grow to confluence, the Hippo kinases are activated and increase the level of phosphorylated YAP in the cytoplasm (Zhao et al., 2007). These findings suggest that the Hippo pathway plays a key role in CIP. However, little is known about how cell density is sensed and transduced to control the subcellular distribution of YAP and TAZ.

The distribution of YAP/TAZ can be regulated by multiple pathways, such as cell surface receptor complexes, apical-basal cell polarity, the actin cytoskeleton, cell-extracellular matrix adhesions, and deformation of nuclear envelopes (Low et al., 2014; Pan, 2010; Panciera et al., 2017; Yu et al., 2015). Because CIP relies on cell-cell communication, cell surface adhesion receptors have been attractive key regulators of YAP and TAZ distribution (Gumbiner and Kim, 2014; McClatchey and Yap, 2012; Schlegelmilch et al., 2011; Silvis et al., 2011). Of the several classes of cell adhesion receptors, much attention has focused on the protein complex comprising E-cadherin and α/β-catenin. Disrupting functional E-cadherin complexes by the removal of extracellular calcium (Schlegelmilch et al., 2011; Varelas et al., 2010), addition of anti-E-cadherin antibodies (Nishioka et al., 2009), or depletion of β-catenin (Kim et al., 2011) results in nuclear retention of YAP and TAZ at high cell density and subsequent loss of CIP. However, a mutant E-cadherin, which comprises a non-functional extracellular domain but retains a normal membrane-tethered cytoplasmic domain with the ability to bind to α/β-catenins, restored CIP in MDCK cells (Benham-Pyle et al., 2015), suggesting that E-cadherin-based cell-cell cohesion *per se* is dispensable for CIP. The identity of E-cadherin-independent cell surface receptors that control the YAP and TAZ distribution during CIP remains unknown. As the cell density increases, cells develop not only the E-cadherin complexes but also other types of cell-cell cohesions including tight junctions, desmosomes, and gap junctions. The formation of these junctions mediates dynamic reorganization of the actin cytoskeleton and intermediate filaments, which causes deformation of cell shapes and cellular organelles, resulting in changes in external mechanical constrain to neighbouring cells. Given that these events are interconnected with each other, the specific contribution of each cell surface receptor in controlling YAP and TAZ distribution during CIP has not been fully understood.

Tight junctions serve not only as a permeability barrier but also as a platform for many signalling pathways, including the Hippo pathway (Matter and Balda, 2003; Zihni et al., 2016). The Hippo kinases, MST-1, MST-2, and LATS1, and YAP and TAZ accumulate at tight junctions as the cell-cell cohesions mature (Zhao et al., 2011). Tight junctions also harbour several tumour suppressor proteins, angiomotin and NF2, which stimulate the phosphorylation of YAP and TAZ (Yi et al., 2011; Zhao et al., 2011). ZO-2 is yet another example of a protein that exhibits a dual localization pattern; like YAP and TAZ, ZO-2 accumulates predominantly in the nucleus at low cell density and in the cytoplasm and at tight junctions at high cell density (Oka et al., 2010). In MDCK cells, ZO-2 associates with YAP and TAZ through its PDZ domain (Dominguez-Calderon et al., 2016; Oka et al., 2010) and facilitates nuclear localization of YAP and TAZ at low cell density (Oka et al., 2010). Additionally, overexpression of ZO-2 has been shown to block cell proliferation, whereas inhibition of ZO-2 induces nuclear retention of YAP/TAZ at high cell density (Dominguez-Calderon et al., 2016). However, the biochemical mechanism by which ZO-2 regulates the distribution of YAP and TAZ remains elusive.

Here, we investigated the function of ZO-2 in relocating YAP from the nucleus to the cytoplasm, as cells grew to confluence and form tight junctions. Independent of the physical constraint of cell shape, apical-basal polarity, and E-cadherin-based cell-cell cohesion, ZO-2 stimulated LATS1-dependent phosphorylation of YAP, thereby promoting the retention of YAP in the cytoplasm in densely cultured cells. ZO-2 was essential for the recruitment of LATS1 to tight junctions as cells grew to confluence. The localization of LATS1 kinase at tight junctions was associated with the accumulation of phosphorylated YAP in the cytoplasm. ZO-2 acted as a scaffold to foster a tripartite complex with LATS1 and YAP, leading to activation of LATS1 by angiomotin and NF2 once the complex was recruited to tight junctions. ZO-2 also inhibited proteasome-mediated degradation of LATS1. These findings delineate a novel mechanism governing CIP, in which ZO-2 utilizes the status of cell-cell cohesion to control the phosphorylation status of YAP by LATS1 in the cytoplasm.

## Results

### External physical constraint is not essential for nucleus-to-cytoplasm translocation of YAP at high cell density

As cell density increases, the shape of cellular architectures changes because of physical constraints from neighbouring cells, together with the establishment of apical-basal polarity, cell-cell cohesions, and reorganization of cellular cytoskeletons. To determine the contribution of external physical constraints on the translocation of YAP from the nucleus to the cytoplasm as cells grow to confluence, we prepared two distinct conditions of MDCK cells at high cell density (Fig. 1 A). In one condition, cells grew to reach confluence over 3 days, permitting them to fully polarize gp135/podocalyxin at the apical cortex and E-cadherin at the lateral cell-cell junctions (Fig. 1, B and C). In the second condition, cells grew to confluence within 24 hours of seeding, which only permitted the establishment of premature apical-basal polarization and weak accumulation of E-cadherin at the cell-cell junctions (Fig. 1, B and C). YAP was exclusively localized to nuclei in cells grown under sparse cell density condition (Fig. 1, B-D). At high cell density, YAP was predominantly localized to the cytoplasm in fully-polarized cells, but was seen both in the cytoplasm and in the nucleus in partially polarized cells (Fig. 1b-d). Consistent with the distribution of YAP, the amount of phosphorylated YAP (at the S127 residue) and LATS1 (at the S909 residue) in the cytoplasm were higher in fully polarized cells than in partially polarized cells (Fig. 1, E and F). By contrast, the phosphorylation of MST-1 and MST-2 (at the T183 and T180 residues, respectively) was unaffected by the degree of cell polarization (Fig. 1 F). Inhibition of LATS1 and YAP did not affect the distribution of gp135 at the apical cortex and E-cadherin at the lateral junctions (Fig. S1), indicating that the Hippo pathway itself was dispensable for the establishment of apical-basal polarity. Collectively, these results suggest that, independent of external physical constraints of cellular architectures, the maturation of cell-cell cohesion is likely to be associated with the activation of LATS1 kinase and the phosphorylation and efficient translocation of YAP from the nucleus to the cytoplasm.

**Figure 1.**
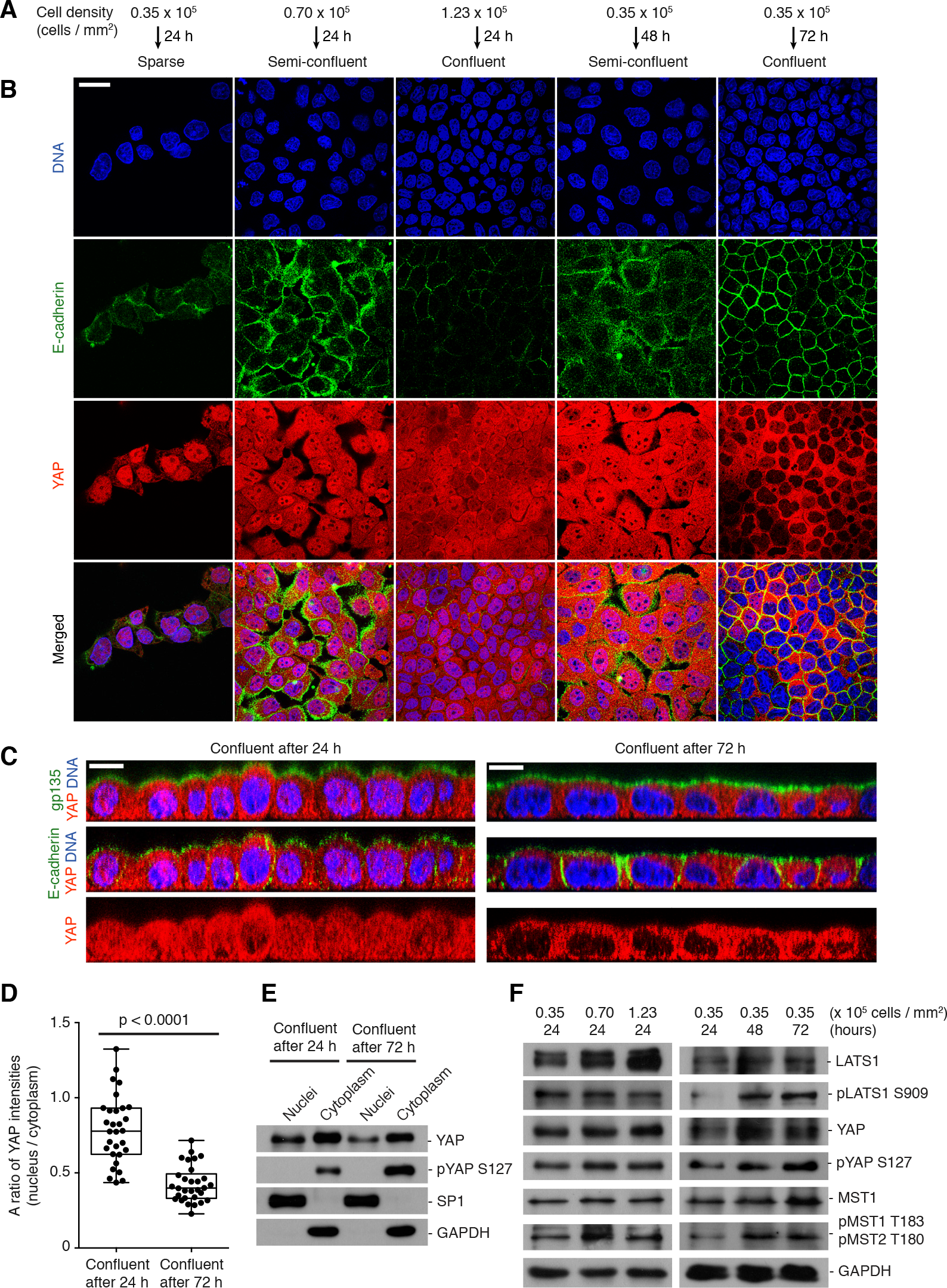
External physical constraint is not essential for nucleus-to-cytoplasm translocation of YAP at high cell density. **(A)** Preparation of MDCK cells under different cell densities and at different stages of apical-basal polarization. Cells were seeded at different cell densities to reach a sparse, semi-confluent, or confluent cell density 24 hours after seeding. Cells were also seeded at distinct cell densities to achieve a semi-confluent or confluent cell density 48 or 72 hours after seeding, respectively. **(B and C)** Representative images of cells at different cell densities and at different stages of apical-basal polarization. Distributions of DNA, E-cadherin, YAP, and gp135 are shown as X-Y views (B) and cross-sectional views (C). Scale bar, 25 µm. **(D)** Quantification of the ratio of YAP intensities in the nucleus relative to that in the cytoplasm in cells at confluent cell density but at different stages of apical-basal polarization. Data in box-and-whisker plots show median (midline), 25^th^ to 75^th^ percentiles (box), and minimum and maximum (whiskers) from n = 30 cells for each condition. *p*-values; student’s t-test. **(E)** Representative immunoblotting images for YAP and S127-phosphorylated YAP (pYAP S127) in the nuclei and the cytoplasmic fraction are shown. SP1 and GAPDH were used as markers for the nuclear fraction and the cytoplasmic fraction, respectively. **(F)** Representative immunoblotting images for LATS1, S909-phosphorylated LATS1 (pLATS1 S909), YAP, S127-phosphrylated YAP (pYAP S127), MST1, T183/T180-phosphorylated MST1/2 (pMST1 T183 / pMST2 T180), and GAPDH from extracts of cells shown in (B). Cell culture conditions (the initial cell densities and the elapsed time after seeding) used to prepare each cell extract are indicated.

### ZO-2 is required for efficient phosphorylation and exclusion of YAP from the nucleus at high cell density

Next, we investigated the role of ZO-2 in the relocation of YAP from the nucleus to the cytoplasm at high cell density. Like YAP, ZO-2 predominantly accumulated in the nucleus at low cell density, and in the cytoplasm and at the tight junctions at high cell density (Fig. 2 A). The translocation kinetics of ZO-2 and YAP were strongly correlated (Fig. 2 A), suggesting a putative role of ZO-2 in exclusion of YAP from the nucleus or in retention of YAP in the cytoplasm. To test these possibilities, we examined the distribution of YAP in ZO-2 depleted cells. Knockdown of ZO-2 using short-hairpin RNA (shRNA) did not alter the retention of YAP in the nucleus at lower cell density (Fig. S2). ZO-2 depleted cells established apical enrichment of gp135 and lateral enrichment of E-cadherin and ZO-1 (Fig. 2 B) but retained a substantial amount of YAP in the nuclei at confluent cell density even three days after seeding (Fig. 2, B-D). In these confluent cells, ZO-2 knockdown did not affect phosphorylation of MST1 and MST2 kinases, but significantly reduced YAP phosphorylation at the S127 residue and marginally reduced LATS1 phosphorylation at the S909 residue (Fig. 2 E). Notably, the amount of LATS1 protein was significantly reduced in ZO-2 depleted cells (Figs. 2 E and 3 A). Expression of ZO-2 transgene restored the levels of LATS1 protein (Fig. 3 B). Based on these results, we conclude that ZO-2 is essential for several events that lead to effective translocation of YAP in fully polarized cells at high density, including the maintenance of the LATS1 protein and the increase in the levels of S909-phosphorylated LATS1 and S127-phosphorylated YAP.

**Figure 2.**
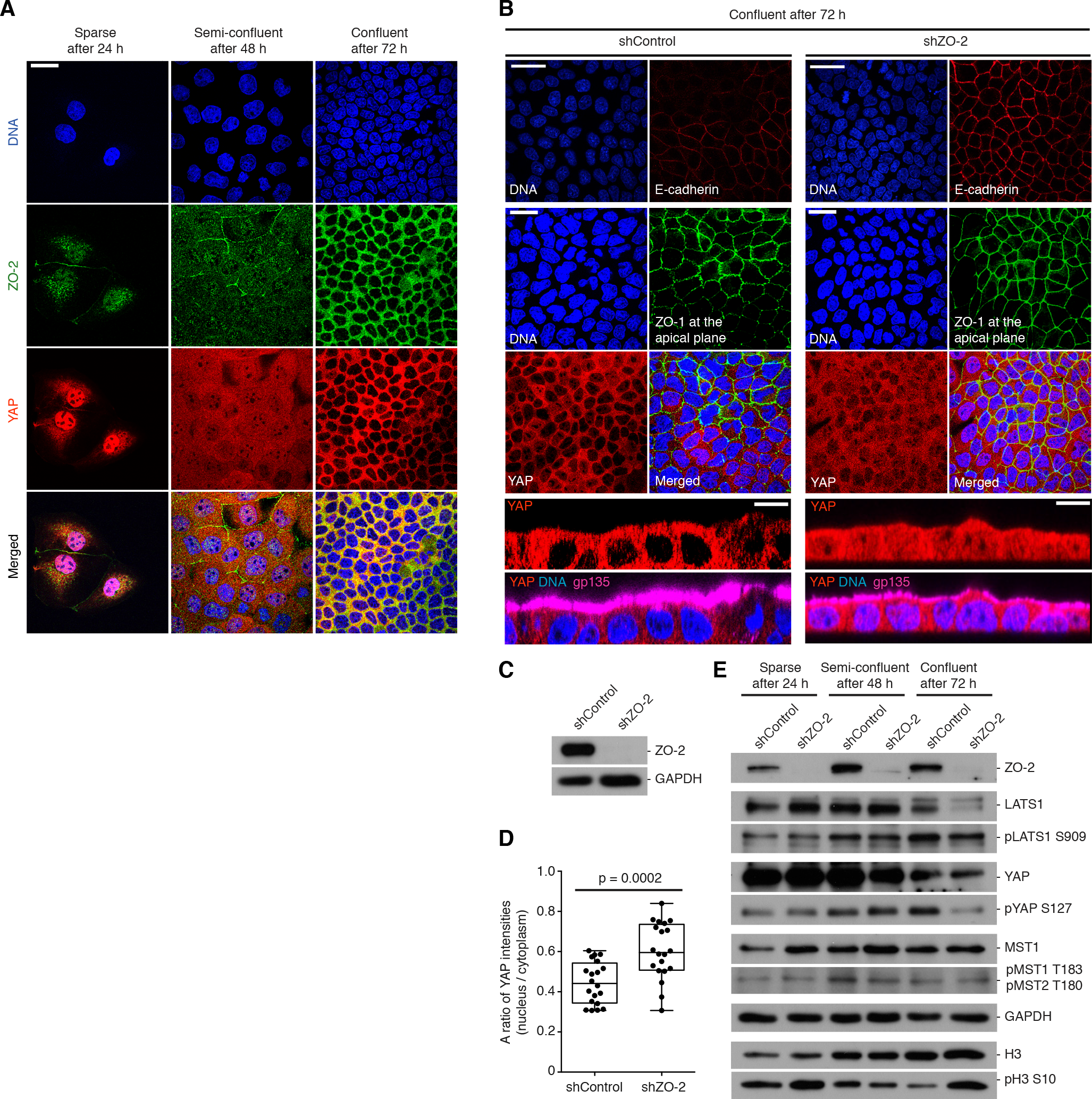
ZO-2 is required for efficient phosphorylation and translocation of YAP at high cell density. **(A)** Representative images of DNA, YAP, and ZO-2 at the lateral plane in MDCK cells under different cell-density conditions. Scale bar, 25 µm. **(B)** Representative images of ZO-2 depleted cells at confluent cell density 72 hours after seeding. Distributions of DNA, E-cadherin, YAP, and ZO-1 are shown as X-Y views (ZO-1 is at the apical plane, and all others at the lateral plane), and those of YAP, DNA, and gp135 are shown as cross-sectional views. Scale bar, 25 µm. **(C)** Representative immunoblotting images for ZO-2 and GAPDH from extracts of ZO-2 depleted cells. **(D)** Quantification of the ratio of YAP intensities in the nucleus relative to that in the cytoplasm in cells shown in (B). Data in box-and-whisker plots show median (midline), 25^th^ to 75^th^ percentiles (box), and minimum and maximum (whiskers) from n = 20 cells for each condition. *p*-values; student’s t-test. **(E)** Representative immunoblotting images for ZO-2, LATS1, S909-phosphorylated LATS1 (pLATS S909), YAP, S127-phosphorylated YAP (pYAP S127), MST1, T183/T180-phosphorylated MST1/2 (pMST1 T183 / pMST2 T180), GAPDH, histone H3 (H3), and S10-phosphorylated histone H3 (pH3 S10) from extracts of control cells and ZO-2 depleted cells at different cell densities.

**Figure 3.**
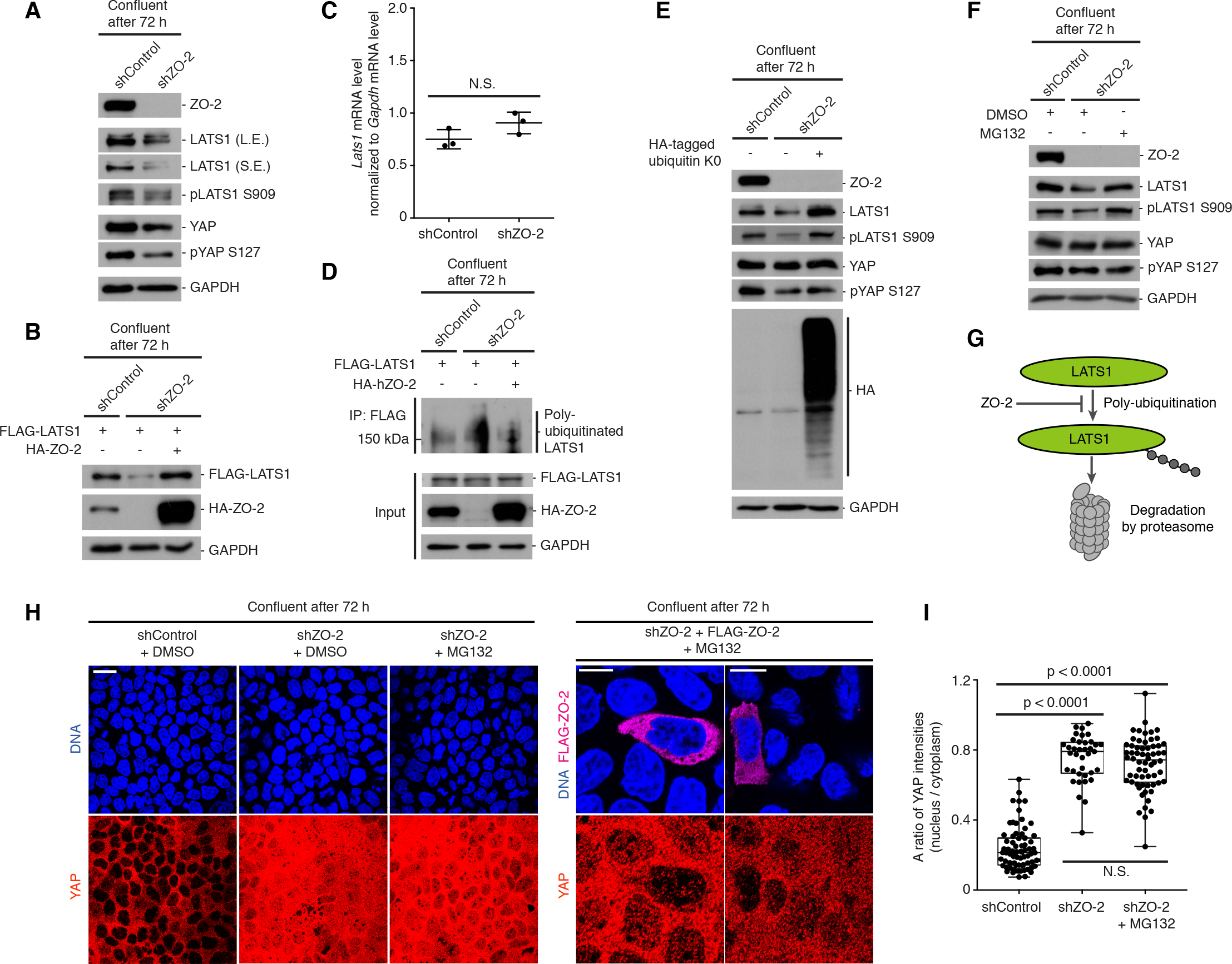
ZO-2 is essential to prevent proteasome-mediated degradation of LATS1 at high cell density. **(A and B)** ZO-2 is required to maintain the levels of LATS1 protein at high cell density. (A) Representative immunoblotting images for ZO-2, endogenous LATS1 (longer exposure: LE, shorter exposure: SE), S909-phosphorylated LATS1 (pLATS S909), YAP, S127-phosphorylated YAP (pYAP S127), and GAPDH from extracts of control and ZO-2 depleted cells. (B) FLAG-LATS1 and HA-ZO-2 immunoblotting images from control and ZO-2 depleted cells with or without ZO-2 transgene are shown. **(C)** *Lats1* mRNA abundance is unaffected by ZO-2 knockdown. The graph depicts the levels of *Lats1* mRNA (normalized to those of *Gapdh* mRNA) measured by RT-qPCR. Plots represent mean values ± s.d. from three independent experiments. *p*-values; Mann-Whitney test. **(D)** ZO-2 limits poly-ubiquitination of LATS1 in cells at high density. Representative immunoblotting images for poly-ubiquitin moiety, FLAG-LATS-1, HA-ZO-2, and GAPDH from the FLAG-LATS1 immuno-precipitates are shown. FLAG-LATS1 were immune-precipitated from extracts of control and ZO-2 depleted cells with or without ZO-2 transgene. **(E)** Inhibition of poly-ubiquitination increases the LATS1 protein level in ZO-2-depleted cells. Representative immunoblotting images for ZO-2, LATS1, S909-phosphorylated LATS1 (pLATS1 S909), YAP, S127-phosphorylated YAP (pYAP S127), and GAPDH from extracts of control cells, and ZO-2 depleted cells with or without the expression of HA-ubiquitin K0 are shown. **(F)** Inhibition of proteasome activity increases the LATS1 protein level in ZO-2 depleted cells. Representative immunoblotting images for ZO-2, LATS1, and S909-phosphorylated LATS1 (pLATS1 S909), YAP, S127-phosphorylated YAP (pYAP S127), and GAPDH from extracts of control cells, and ZO-2 depleted cells with or without MG132 treatment are shown. **(G)** A model of proteasome-mediated degradation of LATS1. ZO-2 antagonizes with poly-ubiquitination and degradation of LATS1. **(H)** Increased levels in LATS1 protein is insufficient for the restoration of nuclear-to-cytoplasm translocation of YAP in ZO-2 depleted cells. Left: representative images of cells depleted of ZO-2 with or without MG132 treatment. Right: representative images of ZO-2 depleted cells with FLAG-ZO-2 transgene after MG132 treatment. The distributions of YAP were visualized 72 hours after seeding. Scale bar, 25 µm. **(I)** Quantification of the ratio of YAP intensities in the nucleus relative to that in the cytoplasm from cells shown in (F). Data in box-and-whisker plots show median (midline), 25^th^ to 75^th^ percentiles (box), and minimum and maximum (whiskers) from n = 71, 37, 62 cells for each condition. *p*-values; student’s t-test.

### ZO-2 is essential to prevent proteasome-mediated degradation of LATS1 at high cell density

ZO-2 may stimulate the nucleus-to-cytoplasm translocation of YAP by maintaining the amount of LATS1 protein as cells grow to confluence. To test this possibility, we investigated whether a higher level of LATS1 protein was sufficient to induce the effective translocation of YAP in ZO-2 depleted cells at high density. Real-time quantitative PCR analysis showed that the level of *Lats1* mRNA was unaffected by ZO-2 knockdown (Fig. 3 C). Given that LATS1 can be poly-ubiquitinated by the ubiquitin ligase CRL4^DCAF1^ (Li et al., 2014), we hypothesized that ZO-2 could inhibit ubiquitination and consequently prevent degradation of LATS1 protein. Immunoblotting against the poly-ubiquitination moiety revealed that the immunoprecipitated LATS1 was poly-ubiquitinated at high cell density (Fig. 3 D). The level of poly-ubiquitinated LATS1 in ZO-2 depleted cells was significantly higher than that in control cells (Fig. 3 D) and returned to a normal level upon re-introduction of ZO-2 by a transgene (Fig. 3 D). The reduction in LATS1 protein levels in ZO-2 depleted cells was suppressed either by over-expression of a dominant-negative ubiquitin allele, Ubi-K0 (Fig. 3 E) or by treatment with a proteasome inhibitor, MG132 (Fig. 3 F). Together, these data suggest that ZO-2 is essential for the maintenance of LATS1 protein through preventing it from ubiquitin-dependent degradation by the proteasome (Fig. 3 G). We then tested whether prevention of LATS1 protein degradation could restore the level of YAP phosphorylation and the relocation of YAP in ZO-2 depleted cells at high density. Although treatment with MG132 restored the levels of LATS1 and its phosphorylation at S909, it did not restore the phosphorylation of YAP at the S127 residue and did not change the distribution of YAP in the nucleus and in the cytoplasm in ZO-2 depleted cells (Fig. 3, H and I). These data indicated that the recovery of LATS1 protein levels alone was insufficient for effective phosphorylation of YAP and its redistribution at high cell density.

### The exit of ZO-2 from the nucleus stimulates efficient nuclear exit of YAP

To test the second possibility whereby the nucleus-to-cytoplasm relocation of ZO-2 could stimulate YAP to exit from the nucleus and/or retain it in the cytoplasm, we sought to create a mutant ZO-2, which would consistently localize to the nucleus irrespective of cell density conditions. ZO-2 contains three nuclear export signal (NES) sequences (Fig. S3 A). A truncated form of ZO-2 (1-590 amino acids), which contained three PDZ domains and lacked the second and the third NESs, predominantly localized in the nucleus at both low and high cell densities (Fig. S3 B). Overexpression of ZO-2[1-590 aa] significantly increased the amount of nuclear YAP at high cell density (Fig. S3 B). Given the physical interaction between ZO-2 and YAP (Oka et al., 2010), these results suggest that the relocation of ZO-2 to the cytoplasm and subsequent reduction in nuclear ZO-2 concentration stimulates the translocation of YAP from the nucleus to the cytoplasm.

### High cell density induces the formation of a LATS1–ZO-2–YAP tripartite complex

To test the third possibility whereby ZO-2 could enable LATS1 to effectively interact with and phosphorylate YAP, we examined if ZO-2 enhanced the interaction between LATS1 and YAP in cells at high density. Immunoprecipitation assays revealed that endogenous ZO-2 was associated with both LATS1 and YAP (Fig. 4, A and B). The interactions among LATS1, ZO-2, and YAP were further enhanced as the cells grew to confluence (Fig. 4, A and B). Depletion of ZO-2 significantly reduced an interaction between LATS1 and YAP (Fig. 4, C and F). These results indicate the formation of a tripartite complex comprising LATS1–ZO-2–YAP in cells at high density. A truncated version of ZO-2 where the SH3 domain was deleted (ZO-2[ΔSH3]), was unable to associate with LATS1 (Fig. 4, D and E) and did not stimulate the interaction between LATS1 and YAP (Fig. 4 F). A mutant form of LATS1 containing mutations in its SH3-domain-binding motifs was unable to bind to ZO-2 (Fig. 4, G and H). LATS1 was detected in FLAG-YAP immunoprecipitates from ZO-2 depleted cells after a transgene that expressed ZO-2 but not ZO-2[ΔSH3] was re-introduced (Fig. 4 F), indicating that an interaction between ZO-2 and LATS1 is essential for LATS1 to efficiently bind to YAP. These results indicate that ZO-2 utilizes its SH3 domain to bind to LATS1 and enhance the interaction between LATS1 and YAP.

**Figure 4.**
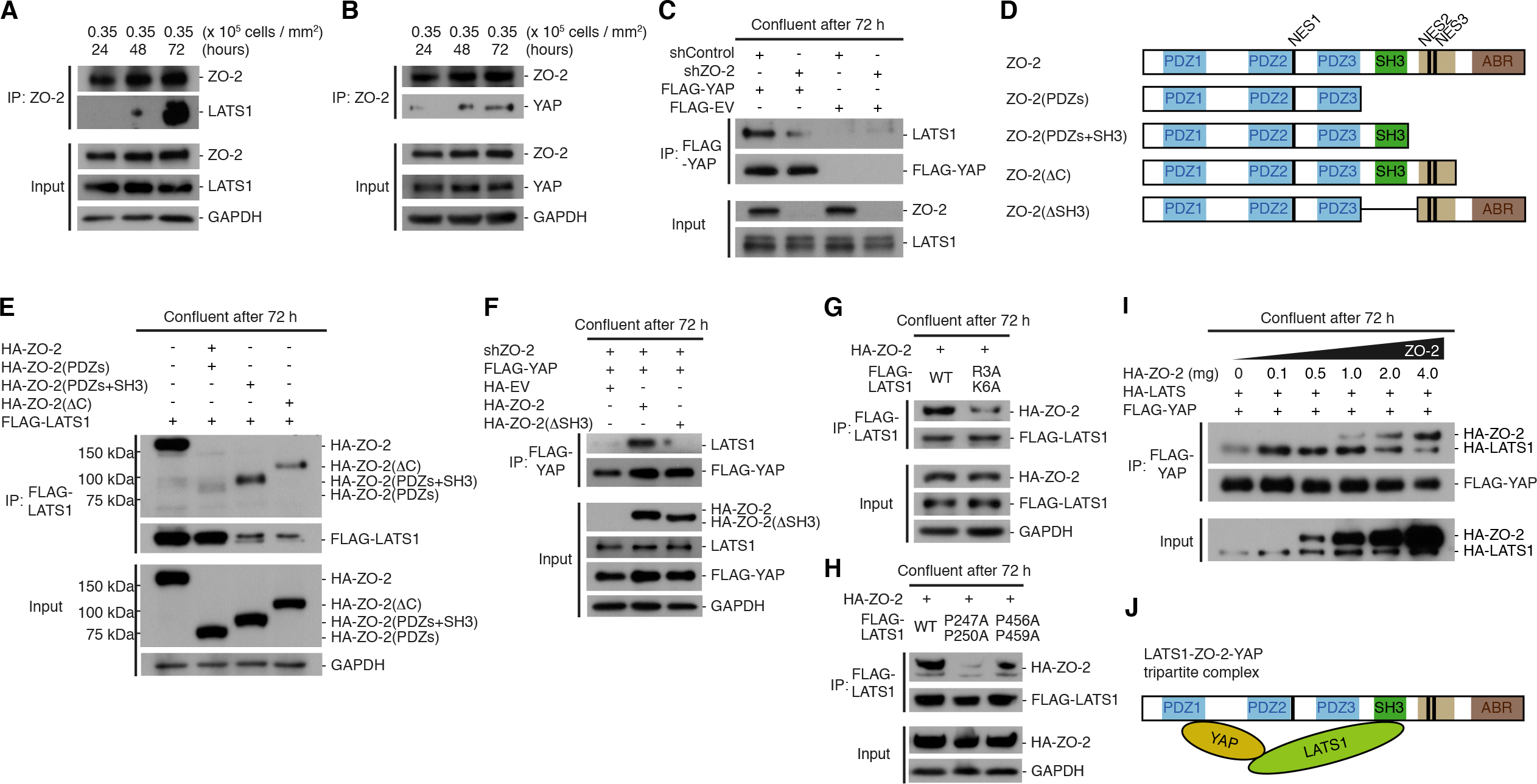
High cell density induces the formation of a LATS1–ZO-2–YAP tripartite complex. **(A and B)** Representative immunoblotting images for (A) ZO-2 and LATS1, (B) ZO-2 and YAP in the ZO-2 immunoprecipitates. ZO-2 was immunoprecipitated from extracts of cells at different cell-density conditions. Levels of ZO-2, LATS1, YAP, and GAPDH in cell extracts are shown as inputs. **(C)** ZO-2 stimulates an interaction between LATS1 and YAP at high cell density. Representative immunoblotting images for LATS1 and FLAG-YAP in the FLAG-YAP immunoprecipitates are shown. FLAG-YAP was immunoprecipitated from extracts of control cells and ZO-2 depleted cells at high density. Levels of ZO-2 and LATS1 in cell extracts are shown as inputs. **(D)** Diagrammatic representation of HA-tagged ZO-2 fragments used in (E) and (F). **(E)** ZO-2 associates with LATS1 via the SH3 domain. Representative immunoblotting images for HA-tagged ZO-2 fragments and FLAG-LATS1 in the FLAG-LATS1 immuno-precipitates are shown. FLAG-LATS1 was immunoprecipitated from the extracts of cells expressing various ZO-2 fragments. Levels of HA-ZO-2 fragments and GAPDH in cell extracts are shown as inputs. **(F)** The interaction between LATS1 and YAP requires the SH3 domain of ZO-2. Representative immunoblotting images for LATS1 and FLAG-YAP in the FLAG-YAP immunoprecipitates. FLAG-YAP was immunoprecipitated from extracts of ZO-2 depleted cells expressing HA-EV, HA-ZO-2, or HA-ZO-2(ΔSH3). Levels of HA-ZO-2, HA-ZO-2(ΔSH3), LATS1, FLAG-YAP, and GAPDH in cell extracts are shown as inputs. **(G and H)** LATS1 interacts with ZO-2 through its SH3 domain-binding motifs. Representative immunoblotting images for HA-ZO-2, FLAG-LATS1, and GAPDH in FLAG-LATS1 immunoprecipitates are shown. FLAG-LATS1 fusions were immunoprecipitated from extracts of cells at high density. The levels of HA-ZO-2, FLAG-LATS1, and GAPDH in cell extracts are shown as inputs. **(I)** ZO-2 acts as a scaffold that associates with both LATS1 and YAP. Representative immunoblotting images for HA-LATS1, HA-ZO-2, and FLAG-YAP in the FLAG-YAP immunoprecipitates. FLAG-YAP was immunoprecipitated from extracts of cells expressing various levels of ZO-2. Levels of HA-ZO-2 and HA-LATS1 in cell extracts are shown as inputs. **(J)** Diagrammatic representation of the formation of a tripartite complex comprising LATS1–ZO-2–YAP.

ZO-2 can interact not only with LATS1 but also with YAP through its first PDZ domain (Oka et al., 2010). We then hypothesized that ZO-2 could serve as a scaffold to facilitate the interaction between LATS1 and YAP. A weak interaction was detected between YAP and LATS1 using FLAG-YAP immunoprecipitates when ZO-2 was severely depleted or at low levels (Fig. 4 I). Increasing levels of ZO-2 led to a significant, corresponding increase in the amount of LATS1 co-immunoprecipitated with FLAG-YAP, and this reached a maximal level with 1mg HA-ZO2, before falling back down to a basal level as HA-ZO-2 concentration was increased (Fig. 4 I). These results imply that an optimal level of ZO-2 is required for the efficient formation of a LATS1–ZO-2–YAP complex. When ZO-2 was highly saturated, a ZO-2–YAP complex (and probably a LATS-1–ZO-2 complex) was predominantly observed at the expense of a LATS1–ZO-2–YAP complex. Based on these biochemical assays, we conclude that ZO-2 serves as a scaffold that enables LATS1 to interact with YAP in a tripartite complex (Fig. 4 J).

### ZO-2 is essential for the recruitment of S909-phosphorylated LATS1 to tight junctions

To further investigate where the LATS1–ZO-2–YAP complex stimulates effective interaction between LATS1 and YAP in cells at high density, we observed the distribution of LATS1 phosphorylated at the S909 residue. Immunostaining revealed the presence of S909-phosphorylated LATS1 in the nucleus at lower cell density and at the apical cell-cell junctions at higher cell density (Fig. 5 A). In ZO-2 depleted cells, this junctional distribution of S909-phosphorylated LATS1 was abolished (Fig. 5 A), and was restored by re-introduction of ZO-2 by a transgene (Fig. 5 B). Consistent with a role of ZO-2 in the recruitment of S909-phosphorylated LATS1 to apical junctions, S909-phosphorylated LATS1 was detected in the ZO-2 immunoprecipitates from cells at high density (Fig. 5 C). To confirm if ZO-2 recruits S909-phosphorylated LATS1 to apically-located tight junctions, we observed S909-phosphorylated LATS1 distribution in cells depleted of ZO-1, another junctional protein essential for the establishment of tight junctions. The junctional enrichment of ZO-2 and S909-phosphorylated LATS1 was abolished in ZO-1 depleted cells (Fig. 5, D and E). Depletion of ZO-1 also reduced the levels of S909-phosphorylated LATS1 and S127-phosphorylated YAP (Fig. 5 F), and led to YAP localization both in the nucleus and in the cytoplasm at high cell density (Fig. 5, G-I). These results suggest that the LATS1–ZO-2–YAP complex enables S909-phosphorylated active LATS1 to interact with and phosphorylate YAP at tight junctions.

**Figure 5.**
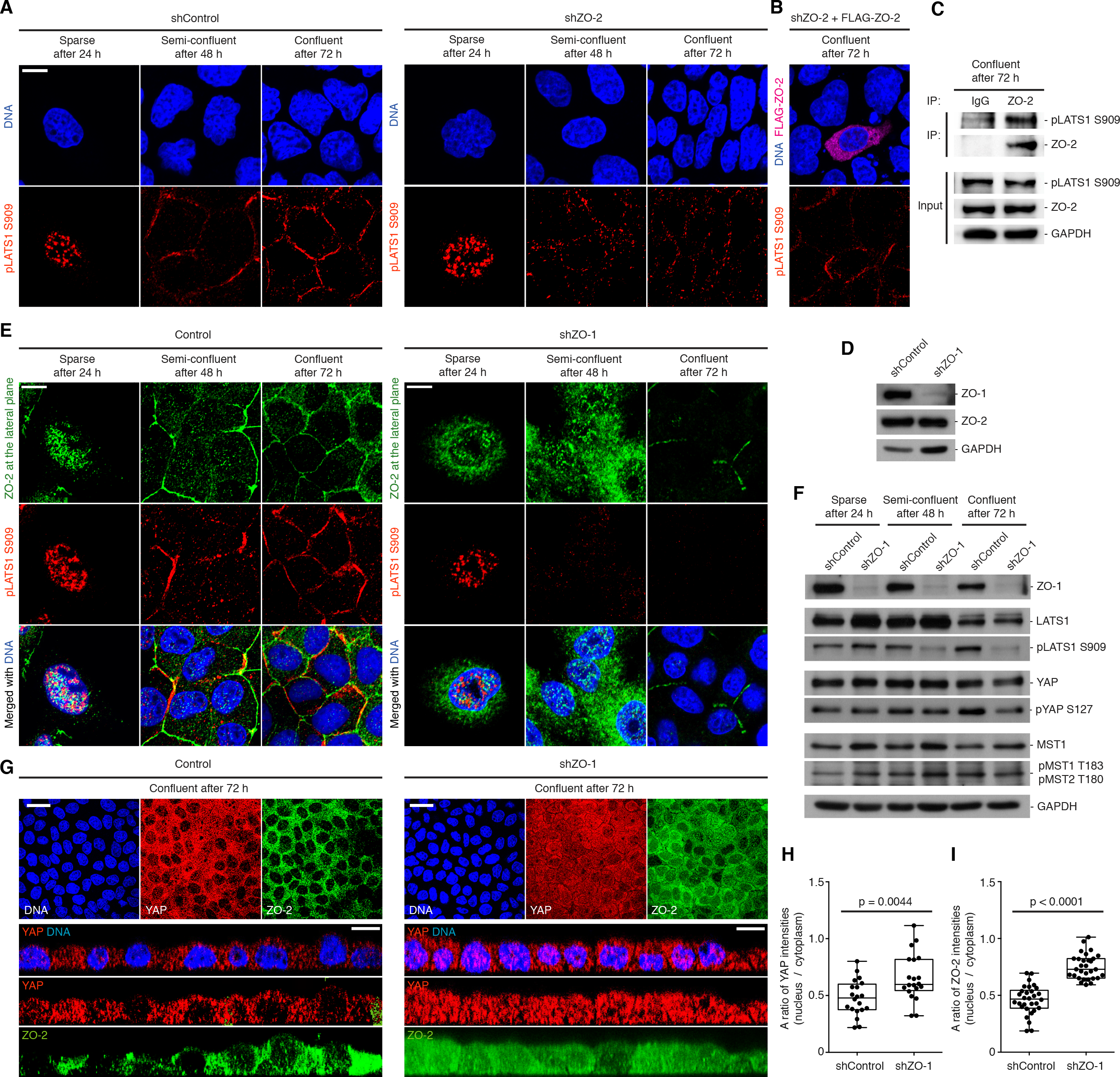
ZO-2 is essential for the recruitment of S909-phosphorylated LATS1 to tight junctions. **(A and B)** ZO-2 recruits S909-phosphorylated LATS1 to the apical junctions. Representative distributions of S909-phosphorylated LATS1 (pLATS1 S909) in control cells and ZO-2 depleted cells (A) and ZO-2 depleted cells with FLAG-ZO-2 transgene (B). Scale bar, 25 µm. **(C)** ZO-2 associates with S909-phosphorylated LATS1. Representative immunoblotting images for S909-phosphorylated LATS1 (pLATS1 S909) and ZO-2 in the ZO-2 immunoprecipitates. ZO-2 was immunoprecipitated from extracts of cells at high density. Levels of pLATS1 S909, ZO-2, and GAPDH in cell extracts are shown as inputs. **(D)** Representative immunoblotting images for ZO-1, ZO-2, and GAPDH in control cells and ZO-1 depleted cells. **(E)** ZO-1-dependent tight junctions are essential for the recruitment of ZO-2 and S909-phosphorylated LATS1 to the apical junctions. Representative distributions of ZO-2 and S909-phosphorylated LATS1 (pLATS1 S909) in control cells and ZO-1 depleted cells. Scale bar, 25 µm. **(F)** ZO-1-dependent tight junction is essential for efficient phosphorylation of LATS1 and YAP. Representative immunoblotting images for ZO-1, LATS1, S909-phosphorylated LATS1 (pLATS1 S909), YAP, S127-phosphorylated YAP (pYAP S127), MST1, T183/T180-phosphorylated MST1/2 (pMST1 T183 / pMST2 T180), and GADPH in extracts of control cells and ZO-1 depleted cells are shown. **(G)** ZO-1-dependent tight junctions are required for effective translocation of YAP from the nucleus to the cytoplasm. Representative distributions of DNA, YAP, and ZO-2 in control cells and ZO-1 depleted cells at high density are shown as X-Y views (top) and cross-sectional views (bottom). Scale bar, 25 µm. **(H and I)** Quantification of the ratio of YAP intensities (H) and ZO-2 intensities (I) in the nucleus relative to that in the cytoplasm from cells shown in (G). Data in box-and-whisker plots show median (midline), 25^th^ to 75^th^ percentiles (box), and minimum and maximum (whiskers) from n = 20 cells (H) and n = 30 cells (I) for each condition. *p*-values; student’s t-test.

### A LATS1–ZO-2–YAP complex is activated by Amot and NF2 at tight junctions for effective phosphorylation and cytoplasmic retention of YAP

Immunoprecipitation assays revealed the association of LATS1 with both YAP and ZO-2 in ZO-1 depleted cells (Fig. 6 A), in which YAP was inefficiently phosphorylated and remained localized to both the nucleus and the cytoplasm (Fig. 5, G and H). This finding raised the possibility that LATS1 in the LATS1–ZO-2–YAP complex could be an inactive state in the cytoplasm and become activated for effective phosphorylation of YAP at tight junctions. No interactions were detected between ZO-1, MST-1, MST-2, and LATS-1 using the FLAG-ZO-1 and FLAG-MST-1 immunoprecipitates (Fig. S4), suggesting that ZO-1 may indirectly stimulate LATS1-mediated phosphorylation of YAP via recruitment of other LATS1 activator(s) to the LATS1–ZO-2–YAP complex at tight junctions (Fig. 6 B). We, therefore, hypothesized that artificial recruitment of other junctional proteins to the LATS1–ZO-2–YAP complex in the cytoplasm could reconstitute protein-protein interactions within the cytoplasmic plaque of tight junctions, thereby restoring the translocation of YAP even in cells without tight junctions, namely ZO-1 depleted cells (Fig. 6 B). Previously, tight junction proteins such as angiomotins (Amot) and NF2, have been identified as regulators of YAP distribution (Yi et al., 2011; Zhao et al., 2011). We used a chemically-inducible dimerization technique (DeRose et al., 2013) to target Amot or NF2 to cytoplasmic ZO-2 in ZO-1 depleted cells (Fig. 6 B). Targeting either FKBP-tagged Amot or FKBP-tagged NF2 to FRB-tagged ZO-2 in the cytoplasm significantly enhanced the relocation of YAP from the nucleus to the cytoplasm within 30 min after addition of a FRB-FKBP dimerization-stimulating reagent, AP21967 (Fig. 6, C and D). In cells co-expressing ZO-2-FRB and either FKBP-Amot or FKBP-NF2, the levels of phosphorylated LATS1 (S909) was lower in the absence of AP21967, but was significantly increased after the addition of AP21967 (Fig. S5). The levels of phosphorylated YAP (S127) showed only marginal increase 30 minutes after AP21967 treatment (Fig. S5). Taking these results together, we conclude that tight junctions serve as a platform for stimulating interactions between the LATS1–ZO-2–YAP complex with angiomotin and NF2, leading to effective activation of LATS1 to phosphorylate YAP and retention of YAP in the cytoplasm.

**Figure 6.**
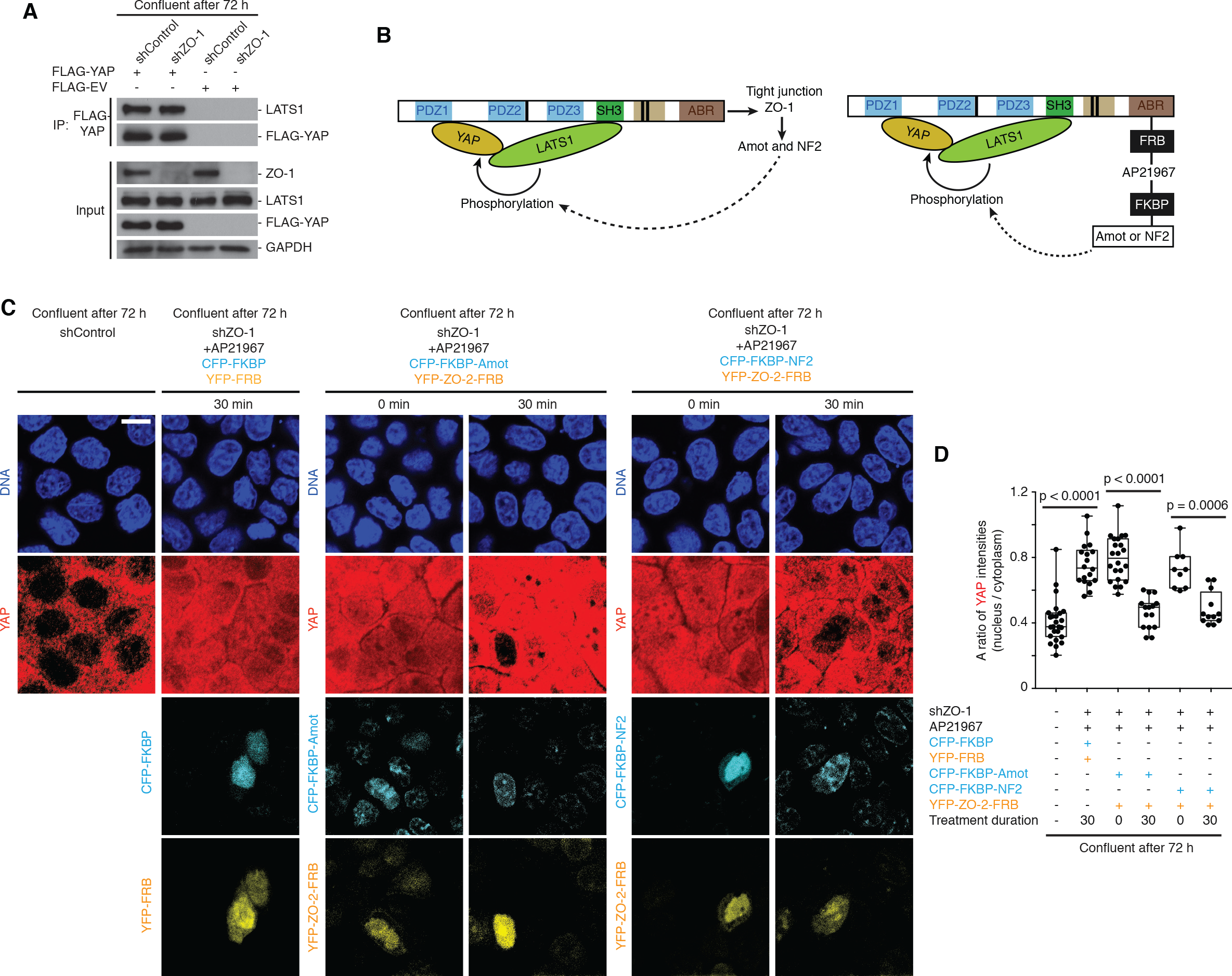
A LATS1–ZO-2–YAP complex is activated by Amot and NF2 for effective phosphorylation and cytoplasmic retention of YAP. **(A)** ZO-1 is dispensable for the formation of LATS1–ZO-2–YAP tripartite complex. Representative immunoblotting images for LATS1 and FLAG-YAP in the FLAG-YAP immunoprecipitates are shown. FLAG-YAP was immunoprecipitated from extracts of control cells and ZO-1 depleted cells at high density. Levels of ZO-1, LATS1, FLAG-YAP, and GAPDH in cell extracts are shown as inputs. **(B-D)** Recruitment of tight-junctional proteins, Amot or NF2, to the LATS1-ZO-2-YAP tripartite complex induces efficient phosphorylation of LATS1 and YAP. (B) Diagrammatic representation of artificial recruitment of Amot and NF2 to ZO-2. (C and D) Recruitment of Amot and NF2 to the LATS1-ZO-2-YAP tripartite complex induces efficient nuclear exit of YAP. (C) Representative images of YAP distribution in ZO-1 depleted cells expressing YFP-FRB and CFP-FKBP after AP21967 treatment and in those expressing YFP-ZO-2-FRB and either CFP-FKBP-Amot or CFP-FKBP-NF2 before and after AP21967 treatment are shown. Scale bar, 25 µm. (D) Quantification of the ratio of YAP intensities in the nucleus relative to that in the cytoplasm from cells shown in (C). Data in box-and-whisker plots show median (midline), 25^th^ to 75^th^ percentiles (box), and minimum and maximum (whiskers) from n = 25, 18, 22, 15, 9, 12 cells for each condition. *p*-values; Mann-Whitney test.

### Discussion

Contact inhibition of proliferation (CIP) has been recognized as a primary mechanism for the regulation of cell and tissue growth, which is controlled by the subcellular distribution of YAP (Gumbiner and Kim, 2014; McClatchey and Yap, 2012). In this study, we presented evidence that tight junctions serve as a platform that transduces the status of cell-cell cohesion into effective phosphorylation and retention of YAP in the cytoplasm. Our findings reveal that the tight junction protein, ZO-2, stimulates LATS1-dependent phosphorylation of YAP in cells at high density in several steps. First, ZO-2 undergoes translocation from the nucleus to the cytoplasm, which facilitates the nuclear exit of YAP and/or the retention of YAP in the cytoplasm. Second, ZO-2 stimulates the interaction between LATS1 and YAP by forming a tripartite complex of LATS1–ZO-2–YAP in the cytoplasm. Third, ZO-2 recruits the tripartite complex to tight junctions, where LATS1 is activated by other junctional proteins, angiomotin and NF2, thereby stimulating the phosphorylation of YAP. Collectively, as cells grow to confluence, tight junctions integrate the formation of a LATS1–ZO-2–YAP complex and the activation of LATS1 by angiomotin and NF2 for effective phosphorylation of YAP. These findings suggest that the cytosolic plaque of tight junctions probably configures these proteins into an ordered structure that facilitates activation of LATS1 kinase by Amot and NF2, as well as promoting a productive interaction between activated LATS1 and YAP for phosphorylation. Our finding of the roles of ZO-2 in linking LATS1 and YAP to tight junctions are consistent with previous observations of physical interaction between YAP and ZO-2 (Oka et al., 2010), ZO-2-mediated modulation of YAP distribution (Dominguez-Calderon et al., 2016), and ZO-2-dependent control of cell cycle progression (Dominguez-Calderon et al., 2016). Although ZO-2 is not essential for the apical enrichment of gp135, ZO-1, or the lateral enrichment of E-cadherin, ZO-2 may also contribute to the reorganization of the actin cytoskeleton at tight junctions, thereby affecting mechanotransduction pathways that will influence the subcellular distribution of YAP. It will be of interest to investigate a potential cross-talk between the ZO-2-dependent control of LATS1 kinase and the reorganization of actomyosin networks at tight junctions.

Further investigation on the mechanism by which the subcellular localization of ZO-2 is coordinated with changes in cell polarization, the integrity of cellular cytoskeletons, and assembly of the apical junctions, will provide deeper insights into the spatio-temporal control of CIP. In particular, how ZO-2 in the nucleus senses the physiological properties of cell density remains unclear. It has recently been reported that mechanical deformation of nuclear envelopes due to physical constraints regulates the nucleus-to-cytoplasm shuttling of a number of proteins including YAP (Elosegui-Artola et al., 2017). Several proteins involved in mechanotransduction between focal adhesions and the extracellular matrix, such as β-catenin (Benham-Pyle et al., 2015) and the LIM domain proteins, zyxin and FHL2 (Nakazawa et al., 2016; Nix and Beckerle, 1997; Smith et al., 2014), also translocate in and out of the nucleus upon mechanical stimulation. ZO-2 may respond to such mechanical deformation of nuclear envelopes or other nucleus-to-cytoplasm shuttling proteins in cells at high density. Other studies have demonstrated that several post-translational modifications of ZO-2, including SUMOylation on K730 (Wetzel et al., 2017), phosphorylation by PKCε on S369 (Chamorro et al., 2009), and O-GlucNAc modification of S257 (Quiros et al., 2013), prevented its entry into the nucleus. It is possible that activities of those enzymes responsible for post-translational modifications of ZO-2 are regulated in a cell density-dependent manner.

We also demonstrated that ZO-2 maintained the level of LATS1 protein during high cell density by protecting it from proteasome-mediated degradation. Our data showed that the SH3 domain of ZO-2 was indispensable for the protection of LATS1, suggesting that a physical interaction between ZO-2 and LATS1 prevented the poly-ubiquitination of LATS1. However, the recovery of LATS1 protein level and its phosphorylation by proteasome inhibition was not sufficient to control cell density-dependent YAP distribution in ZO-2 depleted cells. This results also highlight the importance of ZO-2 for scaffolding active LATS1 and YAP together at tight junctions. LATS kinases are involved not only in the canonical Hippo pathway but also in non-canonical Hippo pathways, such as the control of Ras/Raf signalling (Kilili and Kyriakis, 2010), p53 tumour suppressor (Aylon et al., 2006), and mitotic cell cycle checkpoints (Hergovich and Hemmings, 2012; Visser and Yang, 2010). LATS1 also plays pivotal roles in the Hippo-independent pathways by phosphorylating myosin phosphatase-targeting subunit 1, YPT1 (Chiyoda et al., 2012), 14-3-3γ (Okada et al., 2011), and dual-specificity tyrosine-(Y)-phosphorylation-regulated kinase 1A, DYRK1A (Tschop et al., 2011). The function of ZO-2 in the maintenance of LATS1 protein levels during development and tissue regeneration is an exciting avenue for future exploration.

Tight junctions comprise a vertebrate-specific cell-cell adhesion receptor whose functions and components are evolutionarily conserved (Zihni et al., 2016). In mammals, tight junctions are thought to be analogous to the apically-localizing septate junction complex in *Drosophila* (Tepass et al., 2001). Regulation of YAP distribution by tight junction proteins appears to occur in non-epithelial cells, such as blastomeres in mouse early embryos (Hirate et al., 2013; Lorthongpanich et al., 2013; Nishioka et al., 2009). Hence, tight junctions fulfil multiple functions, by acting as selective permeability barriers and as platforms for intracellular signalling pathways to coordinate cell-density conditions and transcriptional responses. Mechanisms underlying the control of YAP distribution by ZO-2 at tight junctions can be broadly applied to other types of morphogenetic events and the control of organ sizes in animal development, as well as tissue regeneration and the prevention of cancer cell proliferation.

## Acknowledgement

This study was supported by the Mechanobiology Institute of Singapore (to B.C.L.) funded through the National Research Foundation and the Ministry of Education Singapore, the Singapore National Research Foundation (NRF_NRFF2012-08 to F.M.) and the Strategic Japan-Singapore Cooperative Research Program by the Japan Science and Technology Agency and the Singapore Agency for Science, Technology, and Research (1514324022 to F.M.). We are grateful to Andrew Wong and Marius Sudol (Mechanobiology Institute, Singapore) and members in the Low lab for helpful comments on the manuscript.

## Author contributions

The experimental design and presented ideas were developed together by all authors. F.M. and B.C.L guided the study and wrote the manuscript with input from all authors. O.X.L. performed all experiments with a guidance from T.W.C. L.B.L analysed images and performed all statistical analysis.

## Materials and methods

### Plasmids

Plasmids for expression of human LATS1 and MST1 were gifts from Dr. Zhou Dawang (Xiamen University of China). The LATS1 and MST1 cDNAs were sub-cloned into a series of pXJ40 vectors between the NotI and KpnI, and BamHI and XhoI restriction sites, respectively. Human p2xFlag CMV2-ZO-1, p2xFlag CMV2-ZO-2, and p2xFlag CMV2-YAP1-2γ were gifts from Dr. Marius Sudol (National University of Singapore). The ZO-1, ZO-2, and YAP1-2γ cDNAs were sub-cloned into pXJ40 vectors between the XmaI and KpnI, BamHI and KpnI, and NotI and KpnI restriction sites, respectively. pRK5-HA-Ubiquitin-K0 was a gift from Dr. Ted Dawson (Johns Hopkins University). For the chemical inducible dimerization experiments, the ZO-2 and Amot cDNAs were subcloned into YFP-FRB (Addgene *#*20148) and CFP-FKBP (Addgene *#*20160) vectors between XhoI and BamHI restriction sites, respectively. NF2 cDNA was subcloned from a purchased vector (from Addgene *#*32836) into CFP-FKBP vector between EcoRI and BamHI restriction sites. The YFP-FRB and CFP-FKBP Addgene vectors were gifts from Dr. Pakorn Tony Kanchanawong. The primers used in this study for subcloning and mutagenesis are shown in Table S1.

### Cell culture and transfection

MDCK cells were obtained from Dr. David M. Bryant (University of California, San Francisco). Cells were grown in Dulbecco’s Modified Eagles medium (DMEM) with high glucose (Hyclone) supplemented with 10 % fetal bovine serum (Gibco) and 0.2 % sodium bicarbonate (Sigma-Aldrich) at 37 °C in a humidified incubator supplied with 5 % of CO_2_. Cells were transfected with 0.5-5.0 μg plasmids in 125 μL Opti-MEM^®^ medium (Thermo Fisher Scientific) with 7.5 μL of Lipofectamine^®^ 3000 reagent (Invitrogen).

For immunoblotting experiments, cells were seeded in a 35 mm glass-bottom high μ-Dish (ibidi) at three distinct cell densities (0.35 × 10^5^, 0.70 × 10^5^, and 1.23 × 10^5^ cells/cm^2^) to reach sparse, semi-confluent, and confluent culture conditions, respectively, 24 hours after seeding. Alternatively, cells were seeded at a sparse condition (0.35 × 10^5^ cells/cm^2^) and grown for 24, 48, and 72 hours to reach sparse, semi-confluent, and confluent conditions, respectively. The number of cells was measured using LUNA^™^ Automated Cell Counter (Logos Biosystems). For immunofluorescence staining experiments, cells were seeded at three distinct cell densities (0.35 × 10^5^, 0.70 × 10^5^, and 1.23 × 10^5^ cells/cm^2^), and fixed 24 hours post-seeding. Cells were treated with 10 μM MG132 proteasome inhibitor (Sigma Aldrich) or DMSO for 4 hours before fixation.

### Knockdown of ZO-1, ZO-2, and LATS1 by shRNA

Short-hairpin RNAs (shRNAs) targeting ZO-1, ZO-2, and LATS1 were designed using BLOCK-iT^™^ RNAi Designer according to the instructions from the p*Silencer*^™^ siRNA Expression Vectors Instruction Manual. Two single-stranded shRNA template oligonucleotides were heated at 90 °C for 3 minutes and then annealed at 37 °C for 1-2 hours. The annealed shRNA templates were then ligated into either HuSH shRNA GFP cloning vector pGFP-V-RS (TR30007) or HuSH shRNA RFP cloning vector pRFP-C-RS (TR30014). HuSH 29-mer non-effective scrambled shRNA cassettes, pGFP-V-RS (TR30013) and pRFP-C-RS (TR30015), were used as negative controls for pGFP-V-RS and pRFP-C-RS, respectively. Cells were grown with DMEM medium containing 4μ/mL puromycin (Thermo Fisher Scientific) to confluence, seeded into each well of a 96-well plate, and screened for stable cell lines that consistently exhibited a reduction in the level of target proteins. Immunoblotting analyses were used to quantify the levels of protein levels in extracts from cells treated with or without shRNA. The primers used in this study for creating the shRNA constructs are shown in Table S2.

### Cell extract preparation

Transfected cells were harvested, washed twice with PBS, and lysed with RIPA buffer (150 mM of NaCl, 50 mM of Tris-HCl (pH 7.3), 0.25 mM of EDTA, 1 % (w/v) sodium deoxycholate, 1 % (v/v) Triton X-100, 50 mM of sodium fluoride, 1 mM of β-glycerophosphate, 1 mM sodium orthovanadate, and Complete Protease Inhibitor Cocktail Tablets (Roche Applied Science)). Cell lysates were then centrifuged at 21,000 × *g* for 10 minutes to remove insoluble cell debris. The concentration of total protein in each cell lysate was measured using the Pierce^™^ bicinchoninic acid (BCA) Protein Assay Kit (Thermo Fisher Scientific).

### Immunoprecipitation

Cell extracts were centrifuged at 21,000 × *g* for 10 minutes. Supernatants were incubated with primary antibodies overnight at 4 °C, then mixed with Protein A/G PLUS-Agarose (Santa Cruz Biotechnology) for 2 hours at 4 °C. The agarose beads were washed three times with RIPA buffer, and treated with 6X protein SDS loading dye at 85 °C for 6 minutes. FLAG fusion protein was immunoprecipitated with anti-FLAG antibody conjugated with M2 beads (Sigma-Aldrich).

### Immunoblotting

Proteins in the cell extracts and in the immunoprecipitates were separated by SDS-PAGE and then transferred onto a PVDF membrane. The PVDF membrane was incubated with blocking buffer (PBS containing 1 % (w/v) BSA and 0.1 % (v/v) Tween-20) overnight at 4 °C. After incubation with primary antibody in blocking buffer, the membrane was washed three times with washing buffer (PBS, 0.1 % (v/v) Tween-20) at room temperature. Horseradish peroxidase-conjugated anti-mouse antibody (GE Healthcare) was used as a secondary antibody and detected by chemiluminescence using the Pierce^™^ enhanced chemiluminescence Western Blotting Substrate Kit (Thermo Scientific).

### Primary antibodies used for immunoblotting

Rabbit polyclonal anti-ZO-2 antibody (H-110), rabbit polyclonal anti-ZO-1 antibody (H-300), rabbit polyclonal anti-Sp1 antibody (H-225), and mouse monoclonal anti-Ubiquitin antibody (PD41) were purchased from Santa Cruz Biotechnology. Rabbit polyclonal anti-LATS1 antibody (#9153), rabbit polyclonal anti-phospho-LATS1 (Ser909) antibody (#9157), rabbit polyclonal anti-phospho-YAP (Ser127) antibody (#4911), rabbit polyclonal anti-MST1 antibody (#3682), rabbit polyclonal anti-phospho-MST1 (Thr183) and MST2 (Thr180) antibody (#3681), rabbit polyclonal anti-Histone H3 antibody (#9715), rabbit polyclonal anti-phospho-Histone H3 (Ser10) antibody (#9701), and rabbit polyclonal anti-β-actin antibody (#4967) were purchased from Cell Signalling Technology. Rabbit polyclonal anti-FLAG antibody, goat anti-rabbit IgG (A4914), and anti-mouse IgG antibodies conjugated with horseradish peroxidase (A4416) were purchased from Sigma-Aldrich. Rabbit polyclonal anti-HA antibody (#71-5500) was purchased from Invitrogen. Mouse monoclonal anti-GAPDH antibody (AM4300) was purchased from Ambion^®^ Applied Biosystems. Monoclonal mouse anti-E-Cadherin antibody (#610182) was purchased from BD Transduction Laboratories^™^. Rabbit polyclonal anti-YAP antibody was a gift from Dr. Marius Sudol (National University of Singapore).

### Reverse transcription PCR (RT-PCR) and real time quantitative PCR (RT-qPCR)

Cells were seeded at sparsely condition (0.35 × 10^5^ cells/cm^2^) and grown for 72 hours. Cells were subsequently lysed with a buffer RLT (Qiagen) containing 1 % (v/v) β-mercaptoethanol, and then homogenized by passing through a blunt 20-gauge needle fitted to an RNase-free syringe. Total RNA in these cell extracts was purified using RNeasy spin columns (Qiagen). Five micrograms of total RNA were used for first-strand cDNA synthesis with SuperScript^®^ III Reverse Transcriptase Kit (Thermo Fisher Scientific). The first-strand cDNA was then incubated with 0.1 M DTT, RNaseOUT^™^ Recombinant RNase Inhibitor, and SuperScript^™^ III reverse transcriptase at 25 °C for 5 minutes, followed by 50 °C for 60 minutes, and then at 70 °C for 15 minutes. The synthesized cDNA was used as templates for amplification of target genes in RT-PCR and RT-qPCR experiments.

RT-PCR was performed using 100 ng cDNA template, 0.2 mM dNTPs, DyNAzyme II DNA polymerase (Thermo Fisher Scientific), and 10 mM each of the forward and reverse primers. PCR was performed using the following conditions: initial denaturation at 95 °C for 2 minutes, followed by 35 cycles of denaturation at 95 °C for 45 seconds, primer annealing at 55 °C for 45 seconds and elongation at 72 °C for 1 minute, and lastly a final elongation at 72 °C for 10 minutes.

RT-qPCR was performed using 80-140 ng cDNA, SsoFast^™^ EvaGreen^®^ Supermix (Bio-Rad), and 500 nM each of the forward and reverse primers. This reaction mixture was then subjected to an initial denaturation at 95 °C for 30 seconds, followed by 50 cycles of reactions each consisted of denaturation at 95 °C for 5 seconds and combined annealing and extension at 60 °C for 5 seconds. Melt curves were generated at a T_m_ ranging from 65 °C to 95 °C for 5 seconds at each 0.5 °C increment.

### Immunostaining

Immunostaining experiments were performed with cells grown in a 35 mm high glass bottom μ-Dish (ibidi). Cells were washed with PBS, fixed with PBS containing 4 % paraformaldehyde for 15 minutes at 37 °C. Fixed cells were washed with PBS and then with PBS containing 50 mM ammonium chloride. Cells were then permeabilized with PBS containing 0.2 % Triton X-100 for 5 minutes, incubated with blocking buffer (2 % BSA and 7 % FBS in PBS) for 1 hour at room temperature, and incubated with the primary antibodies in blocking buffer overnight at 4 °C. Subsequently, cells were washed five times using PBS buffer containing 0.1 % Triton X-100, and then incubated with the Alexa Fluor^®^ conjugated secondary antibodies and Hoechst 33258 in blocking buffer for 1 hour at room temperature.

### Antibodies used for immunofluorescence staining

Rabbit polyclonal anti-YAP antibody was a gift from Dr. Marius Sudol (National University of Singapore). Rabbit polyclonal anti-phospho-YAP (Ser127) antibody (#4911), rabbit polyclonal anti-LATS1 antibody (#9153), and rabbit polyclonal anti-phospho-LATS1 (Ser909) antibody (#9157) were purchased from Cell Signaling Technology. Rabbit polyclonal anti-ZO-2 (H-110) antibody and goat polyclonal anti-ZO-2 (R-19) were purchased from Santa Cruz Biotechnology. Mouse monoclonal anti-gp135 antibody was obtained from the DSHB (DSHB Hybridoma Product 3F2/D8). Mouse monoclonal anti-FLAG antibody (F3165), rat monoclonal anti-Uvomorulin/E-Cadherin antibody (U3254), and Hoechst 33258 were all purchased from Sigma-Aldrich. Mouse monoclonal anti-HA.11 epitope tag antibody (16B12) was purchased from Covance. Mouse monoclonal anti-Lamin A + C antibody [131C3] (ab8984) was purchased from Abcam. Mouse anti-Occludin antibody (OC-3F10), mouse monoclonal anti-ZO-1 antibody (#33-9100), and all the secondary antibodies conjugated with Alexa Fluor^®^ dyes (405, 488, 568 and 633) (A31556 (Goat anti-rabbit, 405 nm), A31553 (Goat anti-mouse IgG, 405 nm), A11070 (Goat anti-rabbit, 488 nm), A21202 (Donkey anti-mouse, 488 nm), A11036 (Goat anti-rabbit, 568 nm), A21071 (Goat anti-rabbit, 633 nm), A21052 (Goat anti-mouse, 633 nm) were purchased from Invitrogen.

### Ubiquitin Assay

The pRK5-HA-Ubiquitin-K0 plasmid was used for the expression of HA-tagged ubiquitin monomer carrying zero lysines (K0). Two micrograms of the plasmid were transfected in cells treated with shZO-2 48 hours after cell seeding. Cell extracts were prepared as described above.

### Imaging and image processing

Cells were observed at 37 °C with a PlanApochromat 100X 1.4 NA oil immersion lens on a Nikon A1R inverted microscope (Nikon) outfitted with 405 nm, 488 nm, 561 nm, and 640 nm lasers (Coherent Sapphire). The images were taken stepwise at 0.1-0.2 µm per step from the focal plane at the bottom of the cell, through the nucleus, to the top of the cells. The z-stacks were merged and analyzed with using the Volume Viewer function in ImageJ software. The fluorescence intensities of YAP were measured per square µm within the nucleus relative to that in the cytoplasm, and this calculation determines the fold-difference in YAP protein levels between the nucleus and the cytoplasm.

### Statistical Analysis

All statistical tests were performed using GraphPad Prism 7.0. Results presented in graphs represent the mean ± standard error of mean (s.e.m.) or standard deviation (s.d.). The D’Agostino-Pearson omnibus test was used for normality testing of the data. Either the Student’s t-test (two-tailed distribution) or Mann–Whitney test was used to calculate *p*-values. The exact sample numerical value and the statistical methods are indicated in the corresponding figure or figure legend. All experiments with or without quantification were independently repeated at least three times with similar results and the representative data are shown. No statistical method was used to predetermine sample size. The experiments were not randomized. The investigators were not blinded to allocation during experiments or outcome assessment.

### Data availability

All data and materials supporting the findings of this study are available from the corresponding author on reasonable request.

---

**Figure S1.**
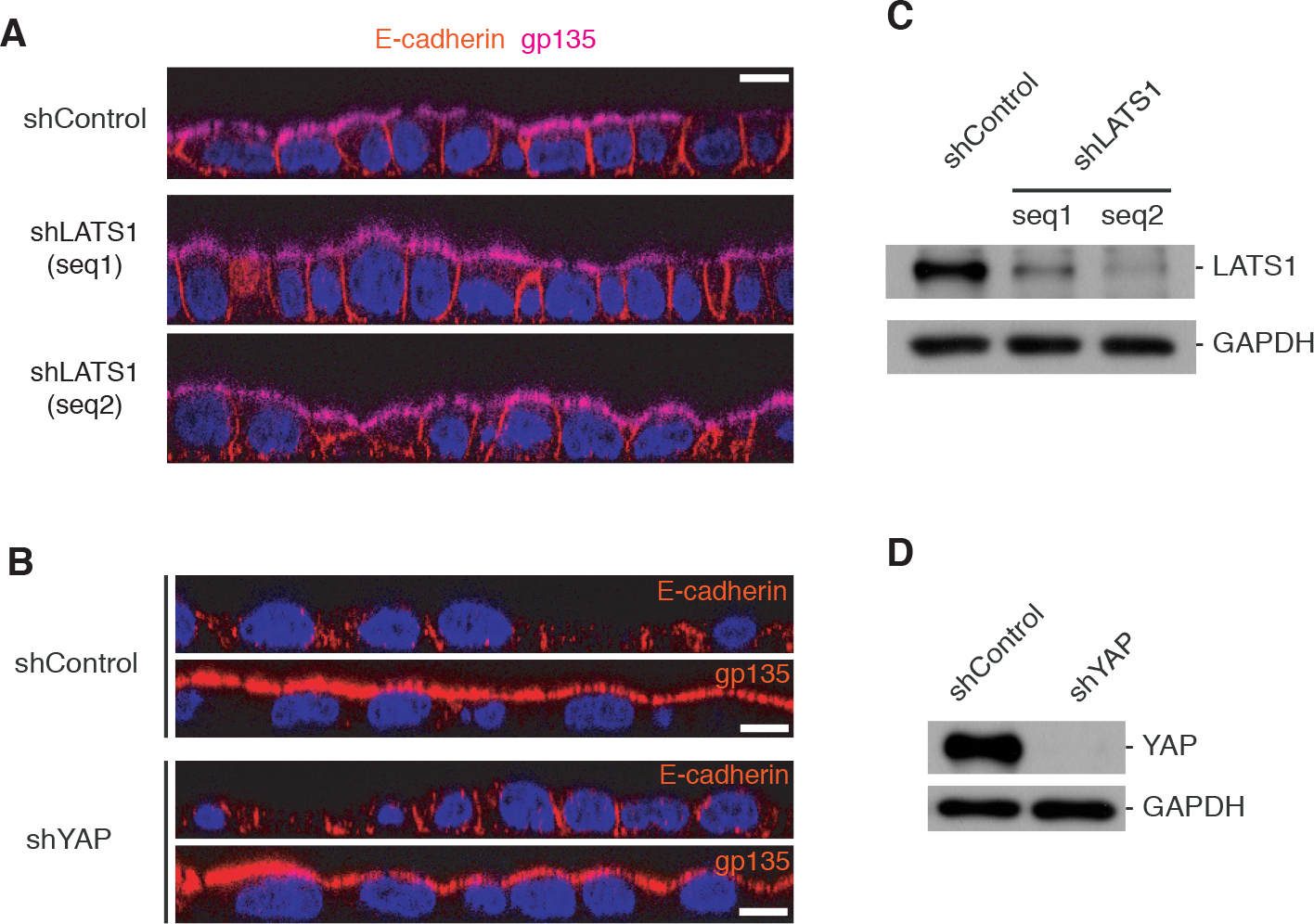
The Hippo pathway is dispensable for the maturation of apical-basal polarity. **(A and B)** Representative images of MDCK cells depleted of LATS1 (A) or YAP (B) at the confluent cell density. The distributions of DNA, E-cadherin, and gp135 are shown as cross-sectional views. Scale bar, 25 µm. **(C and D)** Representative immunoblotting images for LATS1, YAP, and GAPDH from extracts of cells depleted of LATS1 (C) or YAP (D).

**Figure S2.**
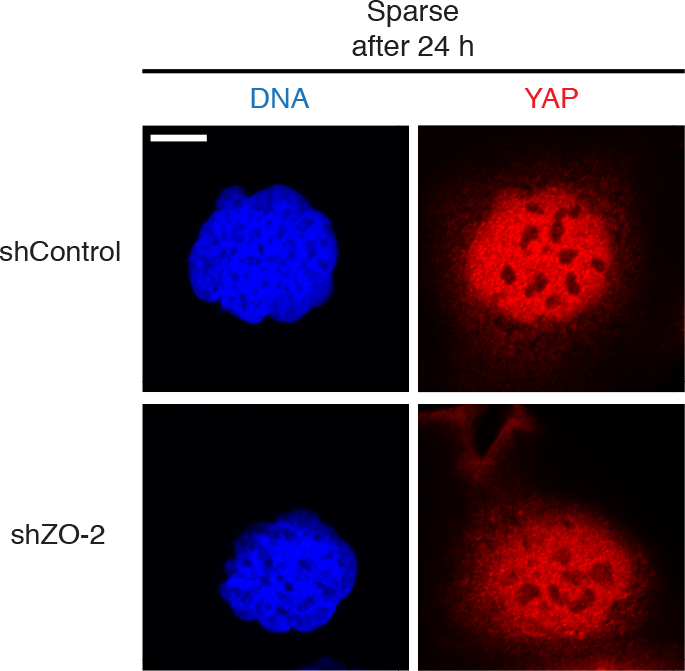
ZO-2 is not essential for YAP to enter the nucleus at low cell density. Representative images of MDCK cells and those depleted of ZO-2 at the low cell density. The distributions of DNA and YAP are shown as X-Y views. Scale bar, 10 µm.

**Figure S3.**
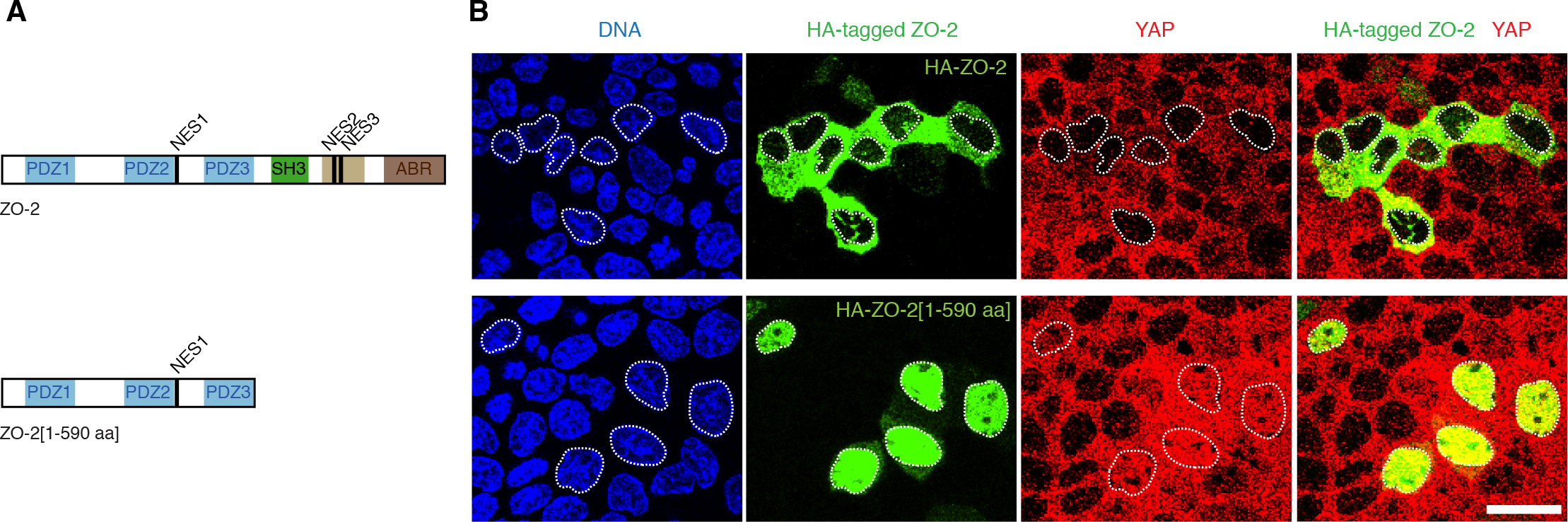
The exit of ZO-2 from the nucleus stimulates efficient nuclear exit of YAP. **(A)** Diagrammatic representation of HA-tagged ZO-2 fragments used in (B). **(B)** Representative images of MDCK cells expressing full-length HA-ZO-2 or HA-ZO-2[1-590 aa]. Distributions of DNA, HA-ZO-2 fusions, and YAP are shown as X-Y views. The area of nuclei in cells expressing HA-tagged ZO2 are marked by white dotted lines. Scale bar, 25 µm.

**Figure S4.**
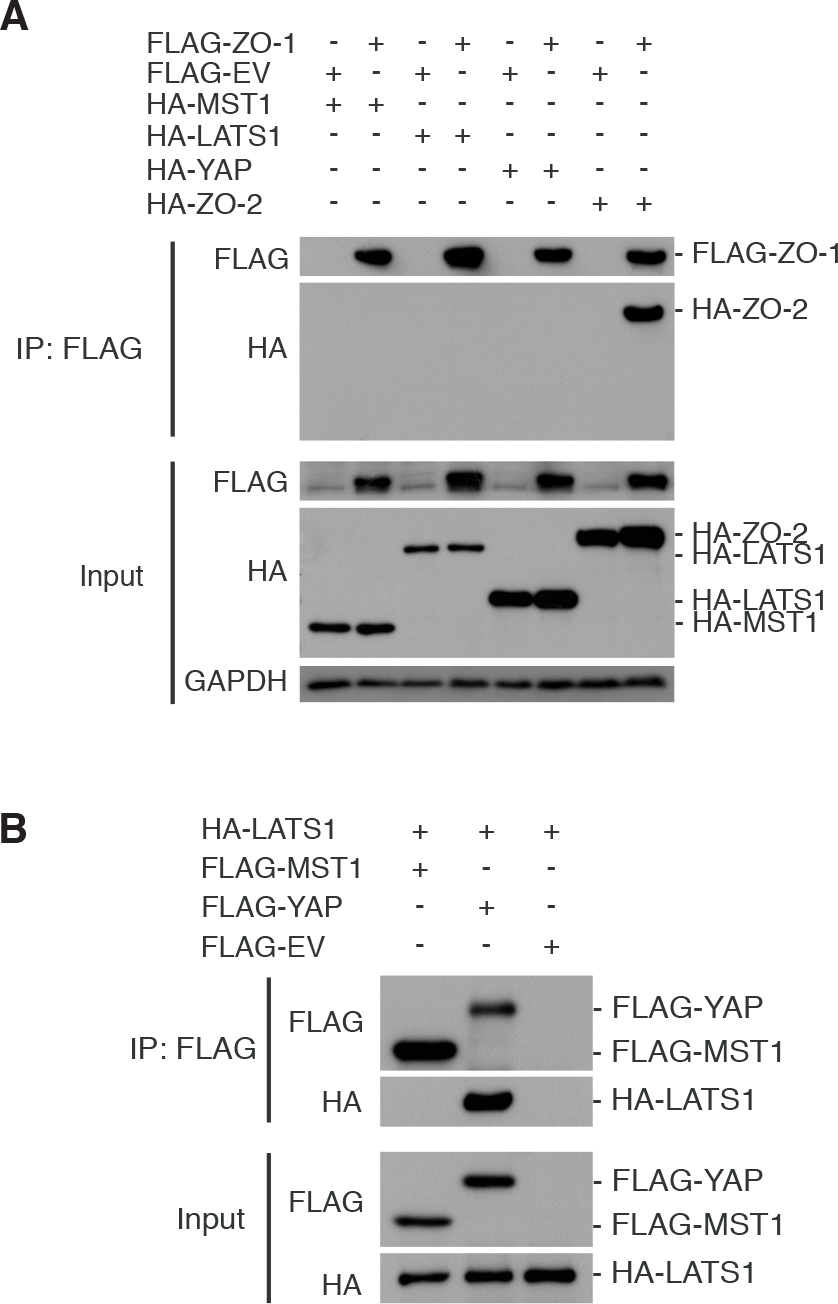
ZO-1 does not directly associate with the Hippo pathway kinases and YAP. **(A)** Representative immunoblotting images for FLAG-ZO-1, HA-ZO-2, HA-MST-1/2, HA-LATS1, and HA-YAP in FLAG-ZO-1 immunoprecipitates. FLAG-ZO-1 was immunoprecipitated from extracts of cells at high density. Levels of FLAG-ZO-1, HA-ZO-2, HA-MST-1/2, HA-LATS1, HA-YAP, and GAPDH from cell extracts are shown as inputs. **(B)** Representative immunoblotting images for FLAG-MST1, FLAG-YAP, and HA-LATS1 in FLAG immunoprecipitates. FLAG fusions were immunoprecipitated from extracts of cells at high density. Levels of FLAG-MST1, FLAG-YAP, HA-LATS1, and GAPDH from cell extracts are shown as inputs.

**Figure S5.**
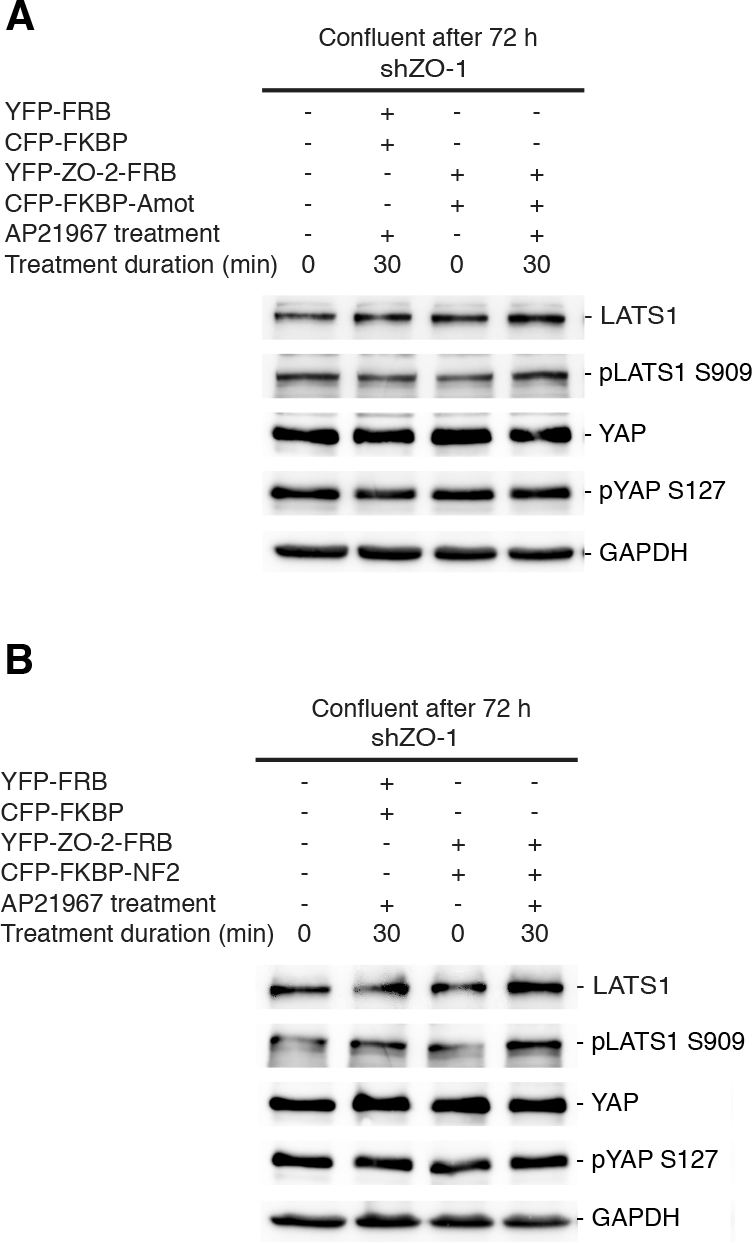
A LATS1–ZO-2–YAP complex is activated by Amot and NF2 for effective phosphorylation and cytoplasmic retention of YAP. **(A)** Representative immunoblotting images for LATS1, S909-phosphoryalted LATS1 (pLATS1 S909), YAP, S127-phosphorylated YAP (pYAP S127), and GAPDH from extracts of ZO-1-depleted cells expressing CFP-FKBP-Amot, CFP-FKBP, YFP-ZO-2-FRB, and YFP-FRB before and after AP21967 treatment are shown. **(B)** Representative immunoblotting images for LATS1, S909-phosphoryalted LATS1 (pLATS1 S909), YAP, S127-phosphorylated YAP (pYAP S127), and GAPDH from extracts of ZO-1-depleted cells expressing CFP-FKBP-NF2, CFP-FKBP, YFP-ZO-2-FRB and YFP-FRB before and after AP21967 treatment are shown.

**Table S1.**
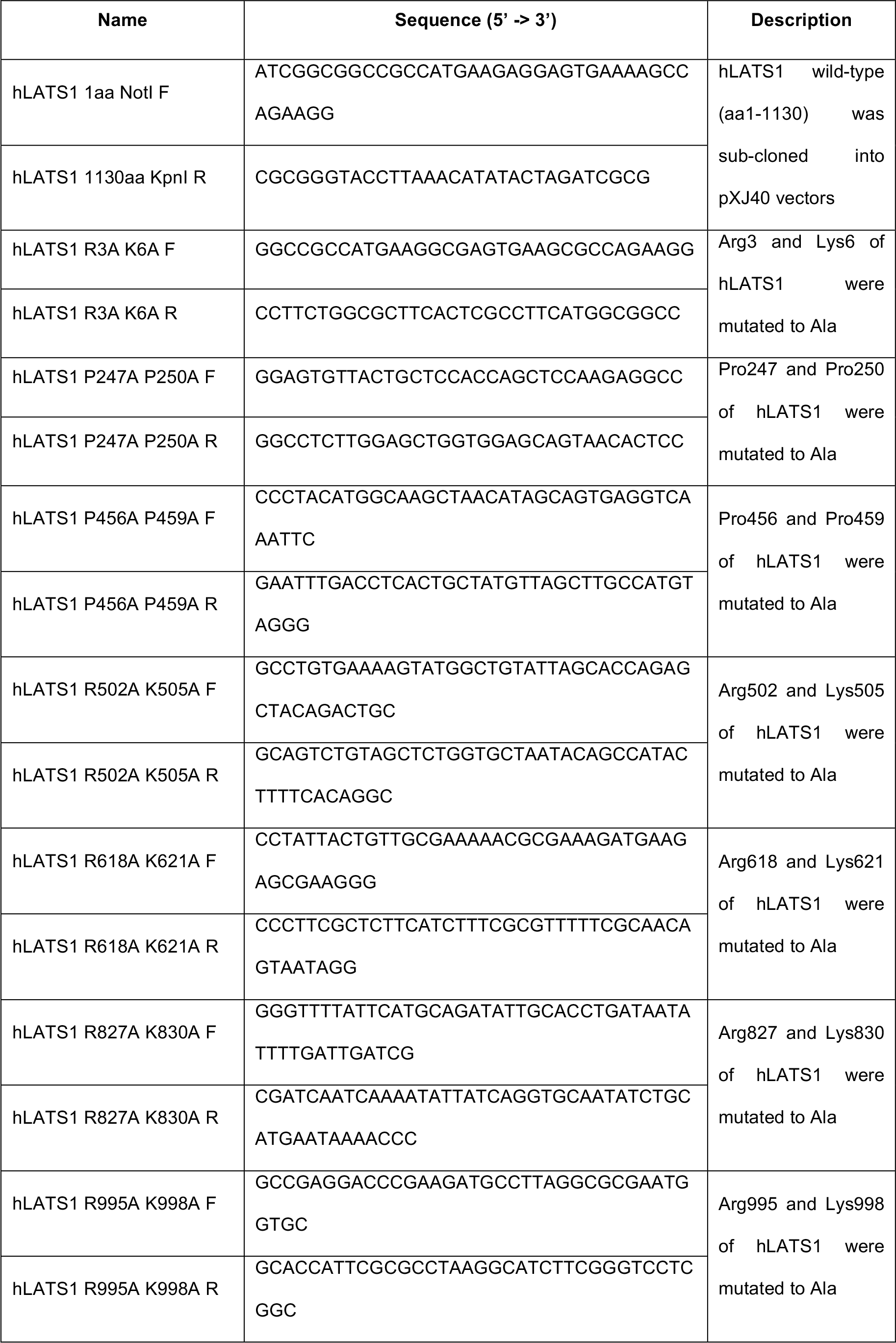
Primers for subcloning and mutagenesis.

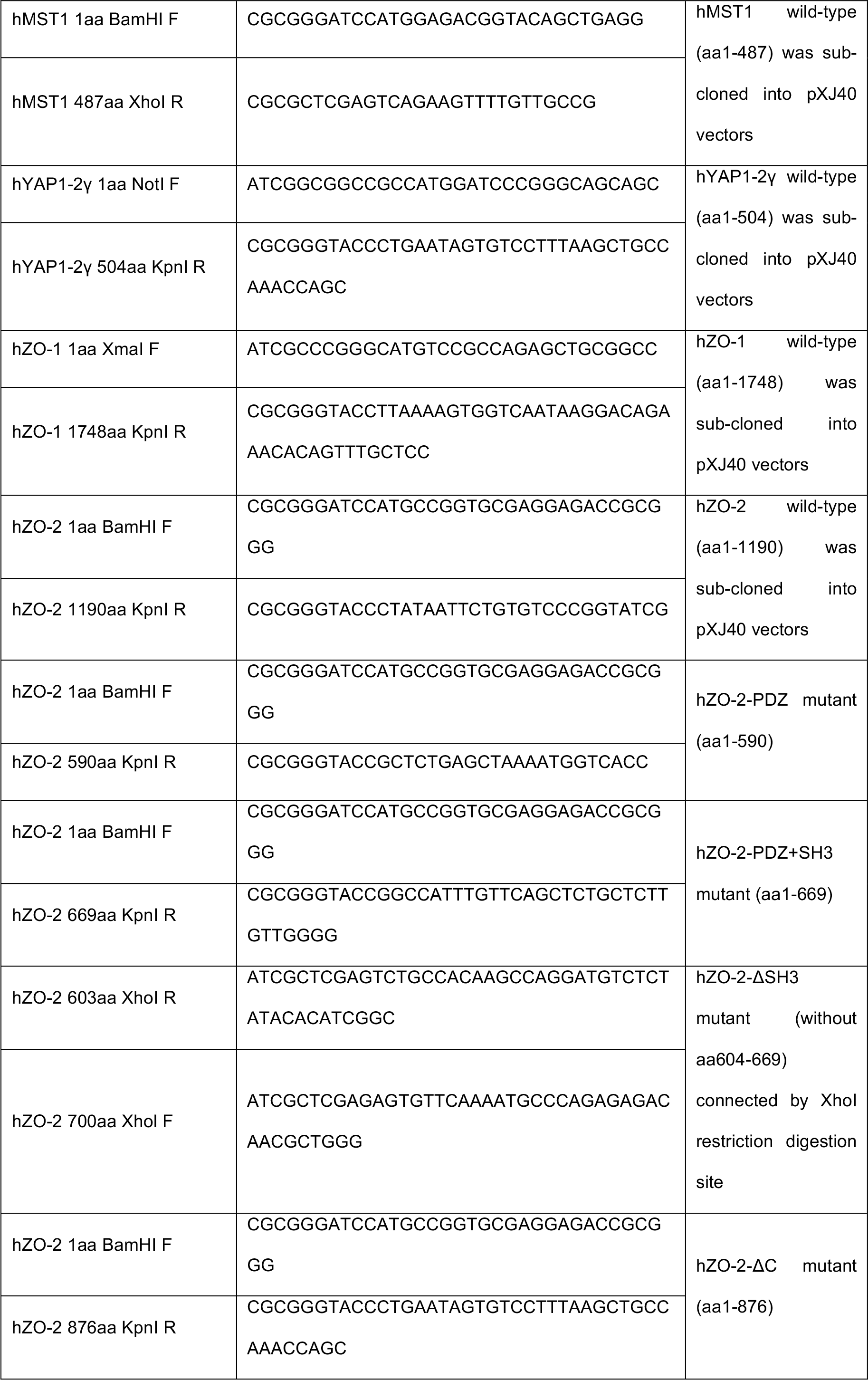

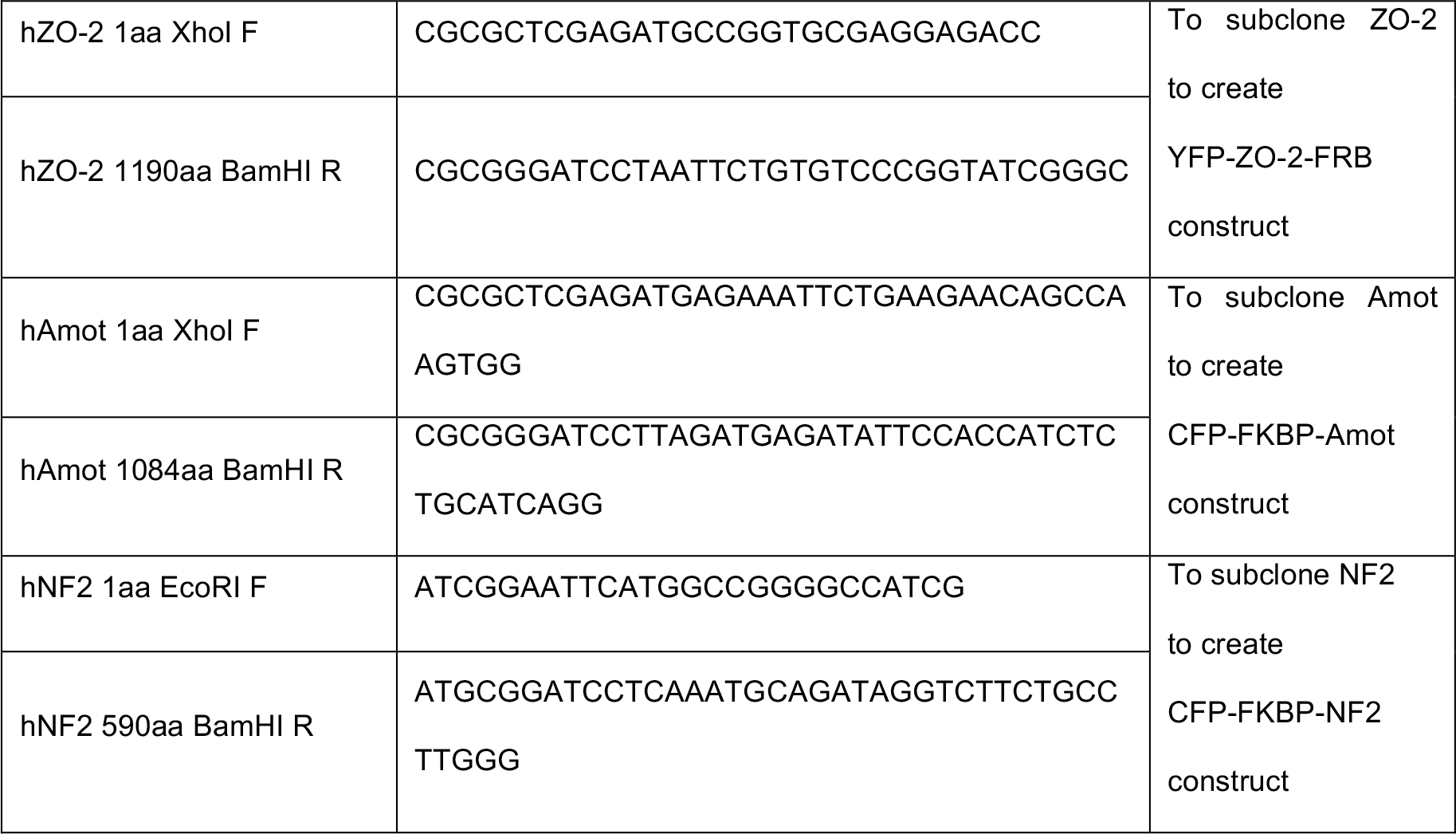

**Table S2.**
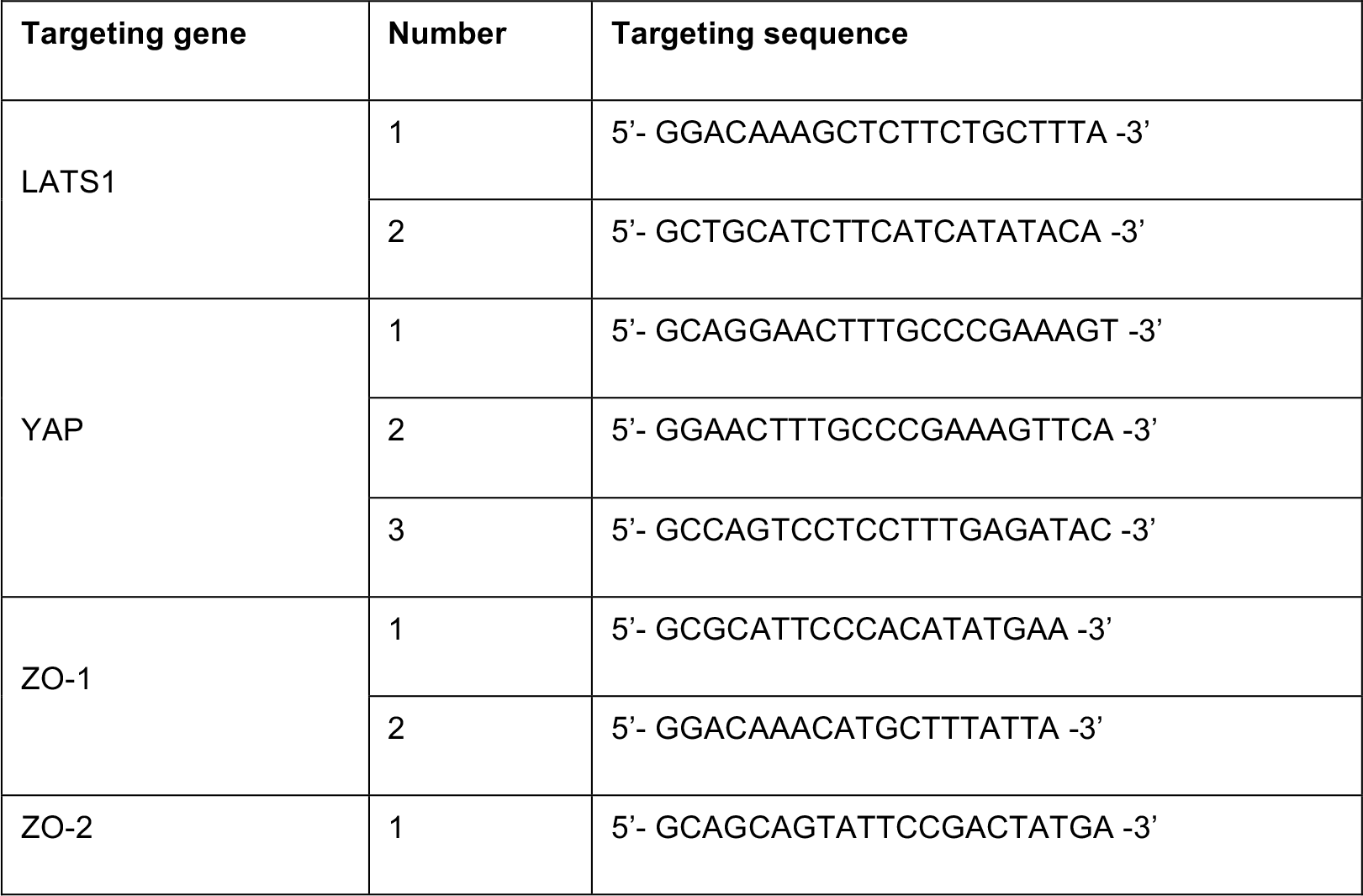
shRNA targeting sequences against LATS1, YAP, ZO-1 and ZO-2 from MDCK cells.

## References

Aylon, Y., D. Michael, A. Shmueli, N. Yabuta, H. Nojima, and M. Oren. 2006. A positive feedback loop between the p53 and Lats2 tumor suppressors prevents tetraploidization. Genes & development. 20:2687–2700.

Benham-Pyle, B.W., B.L. Pruitt, and W.J. Nelson. 2015. Cell adhesion. Mechanical strain induces E-cadherin-dependent Yap1 and beta-catenin activation to drive cell cycle entry. Science. 348:1024–1027.

Callus, B.A., A.M. Verhagen, and D.L. Vaux. 2006. Association of mammalian sterile twenty kinases, Mst1 and Mst2, with hSalvador via C-terminal coiled-coil domains, leads to its stabilization and phosphorylation. FEBS J. 273:4264–4276.

Chamorro, D., L. Alarcon, A. Ponce, R. Tapia, H. Gonzalez-Aguilar, M. Robles-Flores, T. Mejia-Castillo, J. Segovia, Y. Bandala, E. Juaristi, and L. Gonzalez-Mariscal. 2009. Phosphorylation of zona occludens-2 by protein kinase C epsilon regulates its nuclear exportation. Molecular biology of the cell. 20:4120–4129.

Chan, E.H., M. Nousiainen, R.B. Chalamalasetty, A. Schafer, E.A. Nigg, and H.H. Sillje. 2005. The Ste20-like kinase Mst2 activates the human large tumor suppressor kinase Lats1. Oncogene. 24:2076–2086.

Chiyoda, T., N. Sugiyama, T. Shimizu, H. Naoe, Y. Kobayashi, J. Ishizawa, Y. Arima, H. Tsuda, M. Ito, K. Kaibuchi, D. Aoki, Y. Ishihama, H. Saya, and S. Kuninaka. 2012. LATS1/WARTS phosphorylates MYPT1 to counteract PLK1 and regulate mammalian mitotic progression. The Journal of cell biology. 197:625–641.

DeRose, R., T. Miyamoto, and T. Inoue. 2013. Manipulating signaling at will: chemically-inducible dimerization (CID) techniques resolve problems in cell biology. Pflugers Archiv: European journal of physiology. 465:409–417.

Dominguez-Calderon, A., A. Avila-Flores, A. Ponce, E. Lopez-Bayghen, J.V. Calderon-Salinas, J. Luis Reyes, B. Chavez-Munguia, J. Segovia, C. Angulo, L. Ramirez, H. Gallego-Gutierrez, L. Alarcon, D. Martin-Tapia, P. Bautista-Garcia, and L. Gonzalez-Mariscal. 2016. ZO-2 silencing induces renal hypertrophy through a cell cycle mechanism and the activation of YAP and the mTOR pathway. Molecular biology of the cell. 27:1581–1595.

Dong, J., G. Feldmann, J. Huang, S. Wu, N. Zhang, S.A. Comerford, M.F. Gayyed, R.A. Anders, A. Maitra, and D. Pan. 2007. Elucidation of a universal size-control mechanism in Drosophila and mammals. Cell. 130:1120–1133.

Edgar, B.A. 2006. From cell structure to transcription: Hippo forges a new path. Cell. 124:267–273.

Elosegui-Artola, A., I. Andreu, A.E.M. Beedle, A. Lezamiz, M. Uroz, A.J. Kosmalska, R. Oria, J.Z. Kechagia, P. Rico-Lastres, A.L. Le Roux, C.M. Shanahan, X. Trepat, D. Navajas, S. Garcia-Manyes, and P. Roca-Cusachs. 2017. Force Triggers YAP Nuclear Entry by Regulating Transport across Nuclear Pores. Cell. 171:1397–1410e1314.

Galli, G.G., M. Carrara, W.C. Yuan, C. Valdes-Quezada, B. Gurung, B. Pepe-Mooney, T. Zhang, G. Geeven, N.S. Gray, W. de Laat, R.A. Calogero, and F.D. Camargo. 2015. YAP Drives Growth by Controlling Transcriptional Pause Release from Dynamic Enhancers. Molecular cell. 60:328–337.

Gumbiner, B.M., and N.G. Kim. 2014. The Hippo-YAP signaling pathway and contact inhibition of growth. Journal of cell science. 127:709–717.

Hergovich, A., and B.A. Hemmings. 2012. Hippo signalling in the G2/M cell cycle phase: lessons learned from the yeast MEN and SIN pathways. Semin Cell Dev Biol. 23:794–802.

Hirate, Y., S. Hirahara, K. Inoue, A. Suzuki, V.B. Alarcon, K. Akimoto, T. Hirai, T. Hara, M. Adachi, K. Chida, S. Ohno, Y. Marikawa, K. Nakao, A. Shimono, and H. Sasaki. 2013. Polarity-dependent distribution of angiomotin localizes Hippo signaling in preimplantation embryos. Current biology: CB. 23:1181–1194.

Huang, J., S. Wu, J. Barrera, K. Matthews, and D. Pan. 2005. The Hippo signaling pathway coordinately regulates cell proliferation and apoptosis by inactivating Yorkie, the Drosophila Homolog of YAP. Cell. 122:421–434.

Kilili, G.K., and J.M. Kyriakis. 2010. Mammalian Ste20-like kinase (Mst2) indirectly supports Raf-1/ERK pathway activity via maintenance of protein phosphatase-2A catalytic subunit levels and consequent suppression of inhibitory Raf-1 phosphorylation. The Journal of biological chemistry. 285:15076–15087.

Kim, N.G., E. Koh, X. Chen, and B.M. Gumbiner. 2011. E-cadherin mediates contact inhibition of proliferation through Hippo signaling-pathway components. Proceedings of the National Academy of Sciences of the United States of America. 108:11930–11935.

Lei, Q.Y., H. Zhang, B. Zhao, Z.Y. Zha, F. Bai, X.H. Pei, S. Zhao, Y. Xiong, and K.L. Guan. 2008. TAZ promotes cell proliferation and epithelial-mesenchymal transition and is inhibited by the hippo pathway. Molecular and cellular biology. 28:2426–2436.

Li, W., J. Cooper, L. Zhou, C. Yang, H. Erdjument-Bromage, D. Zagzag, M. Snuderl, M. Ladanyi, C.O. Hanemann, P. Zhou, M.A. Karajannis, and F.G. Giancotti. 2014. Merlin/NF2 loss-driven tumorigenesis linked to CRL4(DCAF1)-mediated inhibition of the hippo pathway kinases Lats1 and 2 in the nucleus. Cancer Cell. 26:48–60.

Lorthongpanich, C., D.M. Messerschmidt, S.W. Chan, W. Hong, B.B. Knowles, and D. Solter. 2013. Temporal reduction of LATS kinases in the early preimplantation embryo prevents ICM lineage differentiation. Genes & development. 27:1441–1446.

Low, B.C., C.Q. Pan, G.V. Shivashankar, A. Bershadsky, M. Sudol, and M. Sheetz. 2014. YAP/TAZ as mechanosensors and mechanotransducers in regulating organ size and tumor growth. FEBS letters. 588:2663–2670.

Matter, K., and M.S. Balda. 2003. Signalling to and from tight junctions. Nature reviews. Molecular cell biology. 4:225–236.

McClatchey, A.I., and A.S. Yap. 2012. Contact inhibition (of proliferation) redux. Current opinion in cell biology. 24:685–694.

Nakazawa, N., A.R. Sathe, G.V. Shivashankar, and M.P. Sheetz. 2016. Matrix mechanics controls FHL2 movement to the nucleus to activate p21 expression. Proceedings of the National Academy of Sciences of the United States of America. 113:E6813–E6822.

Nishioka, N., K. Inoue, K. Adachi, H. Kiyonari, M. Ota, A. Ralston, N. Yabuta, S. Hirahara, R.O. Stephenson, N. Ogonuki, R. Makita, H. Kurihara, E.M. Morin-Kensicki, H. Nojima, J. Rossant, K. Nakao, H. Niwa, and H. Sasaki. 2009. The Hippo signaling pathway components Lats and Yap pattern Tead4 activity to distinguish mouse trophectoderm from inner cell mass. Developmental cell. 16:398–410.

Nix, D.A., and M.C. Beckerle. 1997. Nuclear-cytoplasmic shuttling of the focal contact protein, zyxin: a potential mechanism for communication between sites of cell adhesion and the nucleus. The Journal of cell biology. 138:1139–1147.

Oka, T., E. Remue, K. Meerschaert, B. Vanloo, C. Boucherie, D. Gfeller, G.D. Bader, S.S. Sidhu, J. Vandekerckhove, J. Gettemans, and M. Sudol. 2010. Functional complexes between YAP2 and ZO-2 are PDZ domain-dependent, and regulate YAP2 nuclear localization and signalling. Biochem J. 432:461–472.

Okada, N., N. Yabuta, H. Suzuki, Y. Aylon, M. Oren, and H. Nojima. 2011. A novel Chk1/2-Lats2-14-3-3 signaling pathway regulates P-body formation in response to UV damage. Journal of cell science. 124:57–67.

Pan, D. 2010. The hippo signaling pathway in development and cancer. Developmental cell. 19:491–505.

Panciera, T., L. Azzolin, M. Cordenonsi, and S. Piccolo. 2017. Mechanobiology of YAP and TAZ in physiology and disease. Nature reviews. Molecular cell biology. 18:758–770.

Quiros, M., L. Alarcon, A. Ponce, T. Giannakouros, and L. Gonzalez-Mariscal. 2013. The intracellular fate of zonula occludens 2 is regulated by the phosphorylation of SR repeats and the phosphorylation/O-GlcNAcylation of S257. Molecular biology of the cell. 24:2528–2543.

Schlegelmilch, K., M. Mohseni, O. Kirak, J. Pruszak, J.R. Rodriguez, D. Zhou, B.T. Kreger, V. Vasioukhin, J. Avruch, T.R. Brummelkamp, and F.D. Camargo. 2011. Yap1 acts downstream of alpha-catenin to control epidermal proliferation. Cell. 144:782–795.

Silvis, M.R., B.T. Kreger, W.H. Lien, O. Klezovitch, G.M. Rudakova, F.D. Camargo, D.M. Lantz, J.T. Seykora, and V. Vasioukhin. 2011. alpha-catenin is a tumor suppressor that controls cell accumulation by regulating the localization and activity of the transcriptional coactivator Yap1. Sci Signal. 4:ra33.

Smith, M.A., L.M. Hoffman, and M.C. Beckerle. 2014. LIM proteins in actin cytoskeleton mechanoresponse. Trends in cell biology. 24:575–583.

Tepass, U., G. Tanentzapf, R. Ward, and R. Fehon. 2001. Epithelial cell polarity and cell junctions in Drosophila. Annual review of genetics. 35:747–784.

Tschop, K., A.R. Conery, L. Litovchick, J.A. Decaprio, J. Settleman, E. Harlow, and N. Dyson. 2011. A kinase shRNA screen links LATS2 and the pRB tumor suppressor. Genes & development. 25:814–830.

Varelas, X., P. Samavarchi-Tehrani, M. Narimatsu, A. Weiss, K. Cockburn, B.G. Larsen, J. Rossant, and J.L. Wrana. 2010. The Crumbs complex couples cell density sensing to Hippo-dependent control of the TGF-beta-SMAD pathway. Developmental cell. 19:831–844.

Vassilev, A., K.J. Kaneko, H. Shu, Y. Zhao, and M.L. DePamphilis. 2001. TEAD/TEF transcription factors utilize the activation domain of YAP65, a Src/Yes-associated protein localized in the cytoplasm. Genes & development. 15:1229–1241.

Visser, S., and X. Yang. 2010. LATS tumor suppressor: a new governor of cellular homeostasis. Cell cycle. 9:3892–3903.

Wei, X., T. Shimizu, and Z.C. Lai. 2007. Mob as tumor suppressor is activated by Hippo kinase for growth inhibition in Drosophila. The EMBO journal. 26:1772–1781.

Wetzel, F., S. Mittag, M. Cano-Cortina, T. Wagner, O.H. Kramer, R. Niedenthal, L. Gonzalez-Mariscal, and O. Huber. 2017. SUMOylation regulates the intracellular fate of ZO-2. Cell Mol Life Sci. 74:373–392.

Wu, S., J. Huang, J. Dong, and D. Pan. 2003. hippo encodes a Ste-20 family protein kinase that restricts cell proliferation and promotes apoptosis in conjunction with salvador and warts. Cell. 114:445–456.

Wu, S., Y. Liu, Y. Zheng, J. Dong, and D. Pan. 2008. The TEAD/TEF family protein Scalloped mediates transcriptional output of the Hippo growth-regulatory pathway. Developmental cell. 14:388–398.

Yi, C., S. Troutman, D. Fera, A. Stemmer-Rachamimov, J.L. Avila, N. Christian, N.L. Persson, A. Shimono, D.W. Speicher, R. Marmorstein, L. Holmgren, and J.L. Kissil. 2011. A tight junction-associated Merlin-angiomotin complex mediates Merlin’s regulation of mitogenic signaling and tumor suppressive functions. Cancer Cell. 19:527–540.

Yu, F.X., B. Zhao, and K.L. Guan. 2015. Hippo Pathway in Organ Size Control, Tissue Homeostasis, and Cancer. Cell. 163:811–828.

Zanconato, F., M. Forcato, G. Battilana, L. Azzolin, E. Quaranta, B. Bodega, A. Rosato, S. Bicciato, M. Cordenonsi, and S. Piccolo. 2015. Genome-wide association between YAP/TAZ/TEAD and AP-1 at enhancers drives oncogenic growth. Nature cell biology. 17:1218–1227.

Zhao, B., L. Li, Q. Lu, L.H. Wang, C.Y. Liu, Q. Lei, and K.L. Guan. 2011. Angiomotin is a novel Hippo pathway component that inhibits YAP oncoprotein. Genes & development. 25:51–63.

Zhao, B., X. Wei, W. Li, R.S. Udan, Q. Yang, J. Kim, J. Xie, T. Ikenoue, J. Yu, L. Li, P. Zheng, K. Ye, A. Chinnaiyan, G. Halder, Z.C. Lai, and K.L. Guan. 2007. Inactivation of YAP oncoprotein by the Hippo pathway is involved in cell contact inhibition and tissue growth control. Genes & development. 21:2747–2761.

Zihni, C., C. Mills, K. Matter, and M.S. Balda. 2016. Tight junctions: from simple barriers to multifunctional molecular gates. Nature reviews. Molecular cell biology. 17:564–580.

